# Evolutionary adaptation proceeds through a small number of phenotypic modules

**DOI:** 10.1101/2025.09.16.676658

**Authors:** Mohammad Hossein Donyavi, Reza Ghelich, Kara Schmidlin, Grant Kinsler, Kerry Geiler-Samerotte

## Abstract

Understanding how the myriad molecular impacts of mutation percolate to influence higher-order traits and ultimately fitness requires compressing a many-to-many mapping into something tractable. Decades of theoretical work suggest this may be possible because biological systems are modular: effects of perturbation are often funneled through particular pathways or subsystems rather than propagating freely through the organism. Yet few empirical systems have been able to demonstrate such modularity at scale.

Here we show that, even across 774 diverse yeast lineages, fitness variation across 12 drug environments is organized by a strikingly low-dimensional structure defined by only a few inferred phenotypic axes that capture the main patterns of variation. Lineages drawn from multiple evolutionary histories reveal more of these phenotypic axes than those derived from a single selection pressure. Consistent with many of these lineages having evolved under strong selection pressure, their mutations often exhibit broad pleiotropy, affecting nearly all inferred phenotypic axes. However, fitness in any given drug depends on a much sparser subset of the phenotypic modules these axes reflect.

By compressing many-to-many relationships, this low-dimensional framework exposes the modular phenotypic space, as well as the context-dependent contribution of each phenotypic module to fitness that can constrain the pleiotropic effects of adaptive mutations. It also highlights that the apparent complexity of genotype–phenotype–fitness maps depends not only on environmental context but also on the diversity of mutations through which they are observed, laying the groundwork for identifying the key phenotypic modules that matter for fitness.

## 1 Introduction

Predicting how genetic variation shapes phenotype and fitness across diverse contexts remains one of biology’s most enduring challenges, with implications for understanding disease, engineering microbes, and anticipating evolution (Eguchi et al. 2019; Wortel et al. 2022; Zhang et al. 2024; Khodaee et al. 2025; Lässig et al. 2017). This difficulty arises from the complexity of the genotype–phenotype–fitness map, where mutations can be pleiotropic, altering many molecular and physiological phenotypes (Geiler-Samerotte et al. 2020; Kinsler et al. 2020; Paaby and Rockman 2013; Houle et al. 2010). Further, the fitness consequences of these phenotypic changes can vary across environments (Kinsler et al. 2020; Schmidlin et al. 2024; Chen et al. 2023). These many-to-many mappings, linking each mutation to many phenotypes, and each phenotype to fitness in multiple environments, make it difficult to predict fitness effects (Khodaee et al. 2025; Manrubia et al. 2021).

Surprisingly, despite this complexity, recent work shows that variation in mutant fitness across environments can often be captured by low-dimensional models (Kinsler et al. 2020; Otwinowski et al. 2018; Ghosh et al. 2025; Ardell and Kryazhimskiy 2021; Petti et al. 2023). Further, a long history of work has used dimensionality reduction approaches to uncover structure in complex biological systems, including population genetic variation, gene expression data, and quantitative trait architectures (e.g.(Patterson et al. 2006; Novembre et al. 2008; Alter et al. 2000; Lande 1979; Schluter 1996)). In these approaches, latent axes capture shared structure across high-dimensional data; in our setting, these axes correspond to shared patterns of mutant responses across environments. The general success of dimensionality reduction approaches implies an underlying structure in the genotype–phenotype–fitness map, sometimes described in terms of modularity and sparsity (Figure 1).

**Figure 1:**
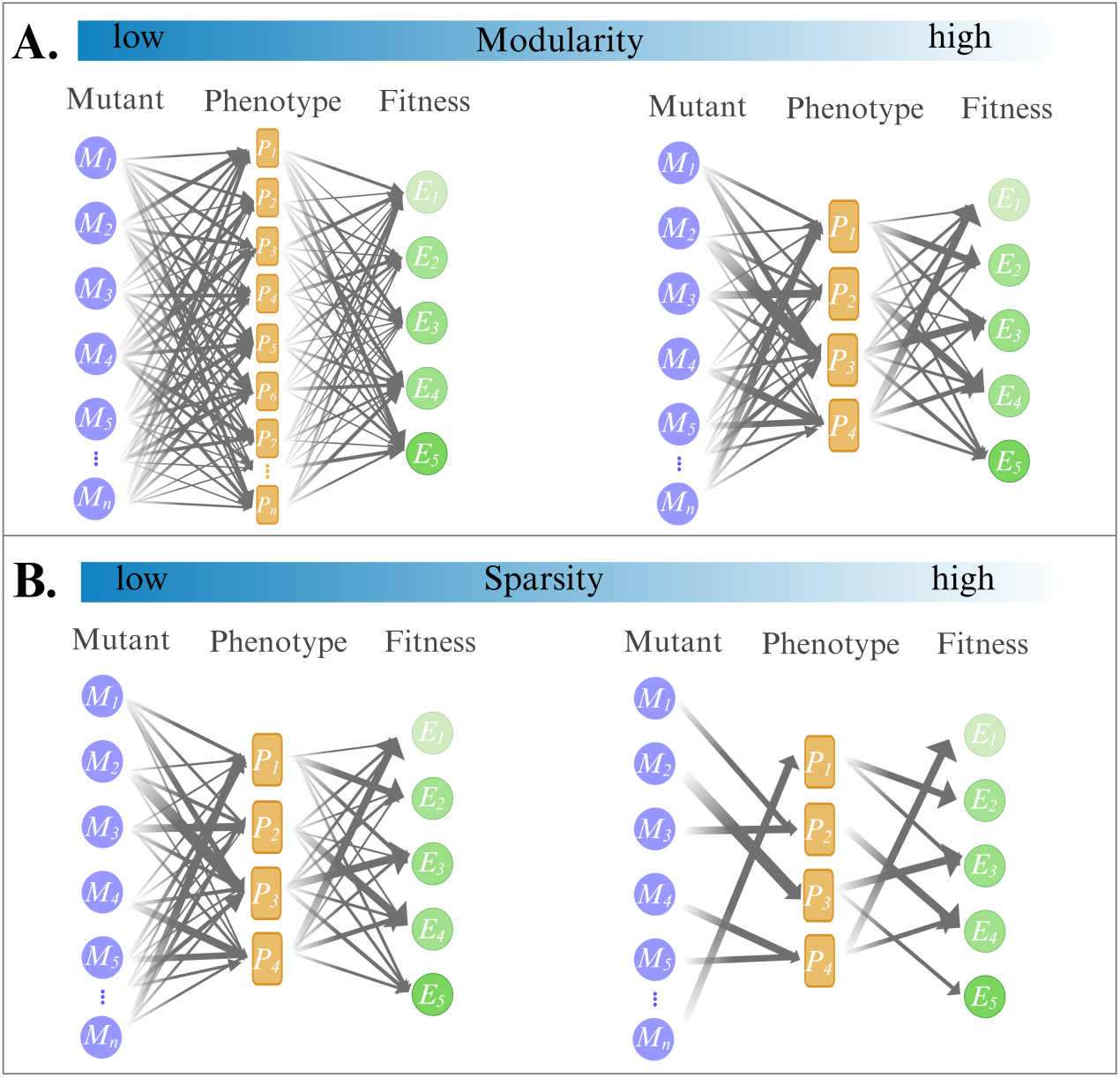
A toy model of the genotype-phenotype-fitness (GPF) map that highlights two key features: modularity and sparsity. These GPF map models are modified from previous work (Kinsler et al. 2020; Petti et al. 2023) to highlight focal concepts in our study. Each map consists of different mutants (blue circles) that influence one or more orthogonal phenotypic modules (yellow squares) to differing degrees (thickness of grey arrows). The phenotypes each affect fitness in one or more environments (green circles) to differing degrees. **(A)** We use the term “modularity” to refer to the number of orthogonal phenotypic axes that mediate fitness (depicted as the number of yellow squares) (Kinsler et al. 2020). In a map with low modularity (top left), the effects of mutations are diffuse, mapping onto a large and complex set of phenotypes that each contribute to fitness in many environments. This represents an architecture where predicting fitness would be more difficult. With high modularity (top right), the effects of different mutations converge on a few phenotypic axes that dictate fitness. This simplifies the map by reducing its effective dimensionality, which could make fitness in unseen environments more predictable. **(B)** We use the term “sparsity” to refer to the density of connections between mutants, phenotypes, and environments (depicted as the number of grey arrows) (Petti et al. 2023). Connections are dense with a low sparsity structure (bottom left). In a high sparsity map (bottom right) a given mutant may only affect a few phenotypes, and a phenotype may only be relevant for fitness in a specific subset of environments. In this study, we first test for modularity, whether fitness variation can be captured by a few shared phenotypic axes, and later assess sparsity, which describes how densely mutations and environments connect to those axes.

Modularity (Figure 1A) refers to the extent to which mutational effects converge on a limited set of independent phenotypic axes that mediate fitness (Geiler-Samerotte et al. 2020; Kinsler et al. 2020; Wagner et al. 2007; Wagner and Altenberg 1996). In a modular system, the myriad consequences of mutation are funneled through a small number of physiological modules, such as stress-response activation or efflux regulation. This concept is closely related to longstanding discussions of phenotypic complexity in evolutionary theory (e.g., (Orr 2000; Martin and Lenormand 2015; Tenaillon et al. 2007; Tenaillon 2014)). Systems with fewer phenotypic modules or dimensions are expected to be more predictable and, in some contexts, less constrained by pleiotropic tradeoffs ((Orr 2000; Tenaillon et al. 2007; Martin and Lenormand 2006)).

Previous work suggests that modularity can exist without requiring mutations to affect only a single phenotypic module (Geiler-Samerotte et al. 2020). Here, we use modularity to refer to the organization of phenotypic variation into a limited number of underlying dimensions, while sparsity describes the extent to which mutational effects are concentrated on those dimensions (Wagner et al. 2007; Petti et al. 2023) (Figure 1B). In this way, the term “sparsity”, as used here, is related to the term “variational modularity” used in previous work (Wagner et al. 2007). Additionally, we and others also use the term “sparsity” to describe whether each phenotypic dimension affects fitness broadly or only in specific contexts (Kinsler et al. 2020; Petti et al. 2023; Ghosh et al. 2025) (Figure 1B). Sparsity improves interpretability because an inferred phenotype is easier to identify when it is influenced by a restricted set of mutations, or contributes to fitness in a restricted set of environments.

The extent of modularity and sparsity in biological systems has been debated for decades (Geiler-Samerotte et al. 2020; Paaby and Rockman 2013; Wagner et al. 2007; Boyle et al. 2017; Hill and Zhang 2012). These properties trace back to Fisher’s early questions about how evolution works, how phenotypes are coupled, which “phenotypic dials” selection can turn independently, and how the structure of the genotype–phenotype map constrains or enables change (Wagner and Altenberg 1996; Draghi and Wagner 2009; Pigliucci 2008; Fisher 1930). Understanding that structure is central not only for explaining evolution, but also for predicting, and ultimately steering, its outcomes (Wortel et al. 2022; Visser and Krug 2014; Nichol et al. 2015; Ogbunugafor 2023).

Drug resistance evolution provides an ideal testbed for probing how modularity and sparsity shape predictability. Mutations that confer resistance to one drug often produce trade-offs in others (Ardell and Kryazhimskiy 2021; Pál et al. 2015; Nichol et al. 2019), and these tradeoffs differ across mutants (Schmidlin et al. 2024; Nichol et al. 2019; Schmidlin et al. 2025). Some mutants show non-monotonic fitness responses to changes in drug concentration, complicating predictions even across similar environments Schmidlin 2024, Schmidlin 2025, Hill 2015, Rosenberg 2018. This variability poses a critical challenge for the design of therapies that seek to trap pathogens in an evolutionary ‘double bind’ by switching to drugs that exploit trade-offs. Given that different resistant mutants possess distinct tradeoffs, it is difficult to design a second-line therapy that eliminates the diverse sub-populations that arise in response to the first drug. But if the apparent diversity in fitness tradeoffs across mutants and drugs conceals modular and sparse structure, evolution may prove more predictable than it seems.

Here we test these ideas by analysing 774 yeast lineages previously evolved in one of twelve antifungal environments Schmidlin2024. Despite extensive genetic diversity, including mutations involving efflux pumps, chaperones, signalling pathways, or membrane components, and despite differing fitness trade-offs and non-monotonic dose responses, variation in fitness across the twelve environments can be captured and predicted by a low-dimensional model. The success of this model indicates strong modularity: diverse mutations converge on a few shared physiological axes that drive fitness across environments. Yet the level of sparsity is asymmetric. The links from mutations to phenotypic modules are not sparse, each mutation perturbs multiple modules, indicating high pleiotropy. On the contrary, the links from phenotypic modules to each environment are sparse. This asymmetry is consistent with expectations from Fisher’s geometric model: adaptation to strong or novel selection proceeds through large, pleiotropic effects, while longer-term stabilizing selection favours modular architectures where phenotypes shape fitness in only a few environments (Fisher 1930; Orr 2000; Wagner and Zhang 2011; Kashtan and Alon 2005; Parter et al. 2007; Kinsler et al. 2024). Our results demonstrate that, even in a system with high pleiotropy, the underlying map has a modular structure that makes evolutionary outcomes more predictable.

## 2 Results

### 2.1 A low-dimensional model predicts the fitness effects of 774 diverse mutations

To explore the extent of modularity in the genotype-phenotype map, we asked whether low dimensional models could make accurate fitness predictions for diverse mutants. While earlier work focused on yeast mutants evolved under a single selection pressure (Kinsler et al. 2020), here we explore yeast evolved under 12 different concentrations or combinations of antifungal drugs (Schmidlin et al. 2024). Like previous work, each evolved mutant lineage we study differs from their shared ancestor by, on average, a single nucleotide mutation (Kinsler et al. 2020). Unlike previous work, the targets of mutation among our 774 lineages are more diverse, including mutations to drug efflux pumps, protein folding chaperones, glucose sensing pathways and cell membrane components (Schmidlin et al. 2024). In addition to diversity at the genotype level, our lineages also have fascinating diversity in their gene-by-environment interactions. Some are broadly drug resistant, others are uniquely resistant or sensitive to narrow drug concentrations, and still others resist single drugs but not their combination (Schmidlin et al. 2024, 2025). Despite this diversity, we found that low dimensional models can predict mutant fitness in drug combinations even when they are built using only single-drug data from a subset of 100 mutants.

Using Singular Value Decomposition (SVD) (Kinsler et al. 2020) (Figure 2A), we built low dimensional models summarizing fitness variation across a training set consisting of 100 randomly drawn mutants in 8 single-drug and no drug conditions. Then we predicted the fitness of the remaining 674 mutants in 4 double-drug combinations (Figure 2B). The fitness predictions we report in figure 2 represent the average across 100 iterations in each of which 100 mutants were randomly chosen for model training. In each iteration, we inferred latent phenotypic dimensions that minimized the mean squared error of the fitness predictions for the held-out 674 lineages. Each latent phenotypic dimension represents an axis of variation in mutant fitness. These dimensions can be thought of as abstract phenotypes inferred from the data, somewhat like principal components inferred from high-dimensional phenotyping studies. We examine the biological interpretation of each inferred phenotypic dimension later. First, we focus on their utility for revealing the structure and modularity of the genotype–phenotype map. In essence, through this modeling, we are asking how well a small number of dimensions can capture how genotype translates to phenotype and fitness.

**Figure 2:**
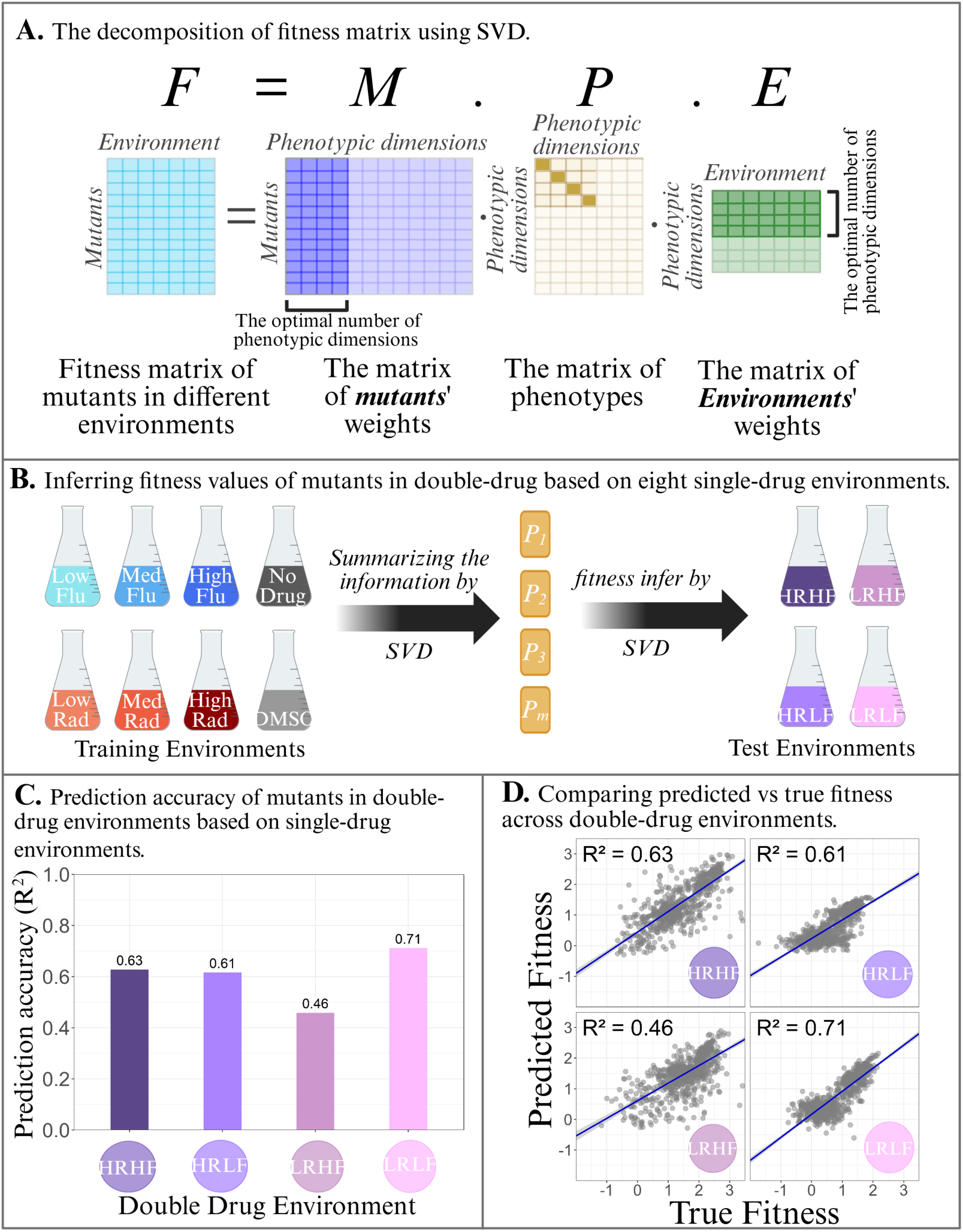
A low-dimensional linear model predicts mutant fitness in double-drug environments from single-drug fitness. **(A)** Schematic of Singular Value Decomposition (SVD) applied to a fitness matrix (F). The matrix is decomposed into mutant weights (M), singular values (Σ), and environmental weights (E), reducing the data into a lower-dimensional space that captures key genotype–phenotype–fitness relationships. **(B)** Training and testing environments. The SVD model was trained on fitness measured in eight single-drug environments (Schmidlin et al. 2024): three Radicicol concentrations (red; 5 *µ*M, 15 *µ*M, 20 *µ*M), three Fluconazole concentrations (blue; 4 *µ*g/ml, 6 *µ*g/ml, 8 *µ*g/ml), and two controls (gray; DMSO, no drug). Model accuracy was tested in four double-drug environments (purple): HRHF (15 *µ*M Rad + 8 *µ*g/ml Flu), HRLF (15 *µ*M Rad + 4 *µ*g/ml Flu), LRHF (5 *µ*M Rad + 8 *µ*g/ml Flu), and LRLF (5 *µ*M Rad + 4 *µ*g/ml Flu). **(C)** Prediction accuracy of the SVD model in the four double-drug environments. **(D)** Scatter plots underlying the *R*^2^ values in panel D, comparing predicted and measured fitness for each of 774 mutants. Predictions were generated by repeatedly training on random subsets of 100 mutants and testing on the remainder; each mutant’s reported prediction is the average across all iterations where it was held out of training.

The median number of dimensions selected across iterations, based on the lowest mean squared error, was 4 (Figure S1). Twice that many dimensions were required to capture variation in random permutations of our data, suggesting that the low dimensional structure we observe is biologically meaningful (Figure S2, S3). Even though our set of mutants and gene-by-environment interactions is diverse, we found that low dimensional SVD models were effective at predicting fitness of held out mutants in held out environments (Figure 2C–D) and at accurately reconstructing the training data (Figure S4). Prediction accuracy across the four double-drug environments ranged from 46% to 71%, which is comparable to previous studies that used similar approaches to show modularity in genotype–phenotype–fitness relationships (Kinsler et al. 2020; Ghosh et al. 2025). Notably, the accuracy of SVD-based predictions exceeds the correlation between biological replicates (*R*^2^ across replicate pairs ranges from 0.42 to 0.59) (Schmidlin et al. 2024), because the replicates were averaged before performing SVD. SVD-based models also outperform simpler approaches in which fitness in a double-drug environment is predicted by adding or averaging fitness across the corresponding single-drug conditions (Figure S5). The accuracy of our SVD-based model remains high even when predictions are generated after a single random iteration (Figure S6).

This predictive success implies that diverse mutational effects converge on a few shared phenotypic dimensions, the hallmark of modularity introduced in figure 1A. The fact that the main axes of phenotypic variation learned from single-drug data could generalize to drug combinations suggests that fitness variation is structured by a small number of underlying processes that repeatedly shape outcomes across conditions. In other words, despite mutations affecting diverse genes and pathways, from efflux pumps to chaperones and signaling components, their phenotypic consequences appear to converge on a limited set of inferred phenotypic ‘dimensions’ or ‘modules’ that represent major contributors to fitness.

Previous work suggests that the lower prediction accuracy of SVD dimensional models in certain environments may arise because not all phenotypic modules are relevant in every condition (Kinsler et al. 2020; Ghosh et al. 2025). Some key phenotypic effects of mutation remain hidden when the training set lacks the environments where these phenotypes constrain fitness. In this view, only a small subset of phenotypes determines fitness in any given environment, while pleiotropic effects on other phenotypes remain latent until other contexts are considered (Kinsler et al. 2020; Ghosh et al. 2025). Thus, modularity both enables and constrains predictability. The shared phenotypic modules make low-dimensional modeling possible, but context-specific ones define the upper bound on prediction accuracy. In the next section, we test these ideas by asking how prediction accuracy changes as we expand the number of environments, mutants, and model components used for training.

### 2.2 Rapid Saturation of Prediction Accuracy Reveals Modular Structure and Its Limits

To assess how modular structure constrains and enables fitness prediction across environments, we examined how prediction accuracy changes as we systematically increase the number of training environments. For each training set size, we repeated the analysis 100 times, randomly selecting a new subset of environments for training in each iteration. We performed this analysis in our focal dataset (12 environments; Figure 3A) and in a very similar dataset where fitness was measured across more environments (45 environments; Figure 3B) (Kinsler et al. 2020). In both datasets, prediction accuracy increased sharply with the first few added environments before rapidly plateauing (Figure 3C). The number of phenotypic dimensions required for high predictive power showed a similar pattern, leveling off after only a few dimensions (Figure S7). Even across model iterations where the ‘optimal’ number of dimensions (the number that minimized mean squared error) was greater, predictive power does not improve (Figure S7). This rapid saturation suggests that a small, representative subset of environments captures most of the predictable signal.

**Figure 3:**
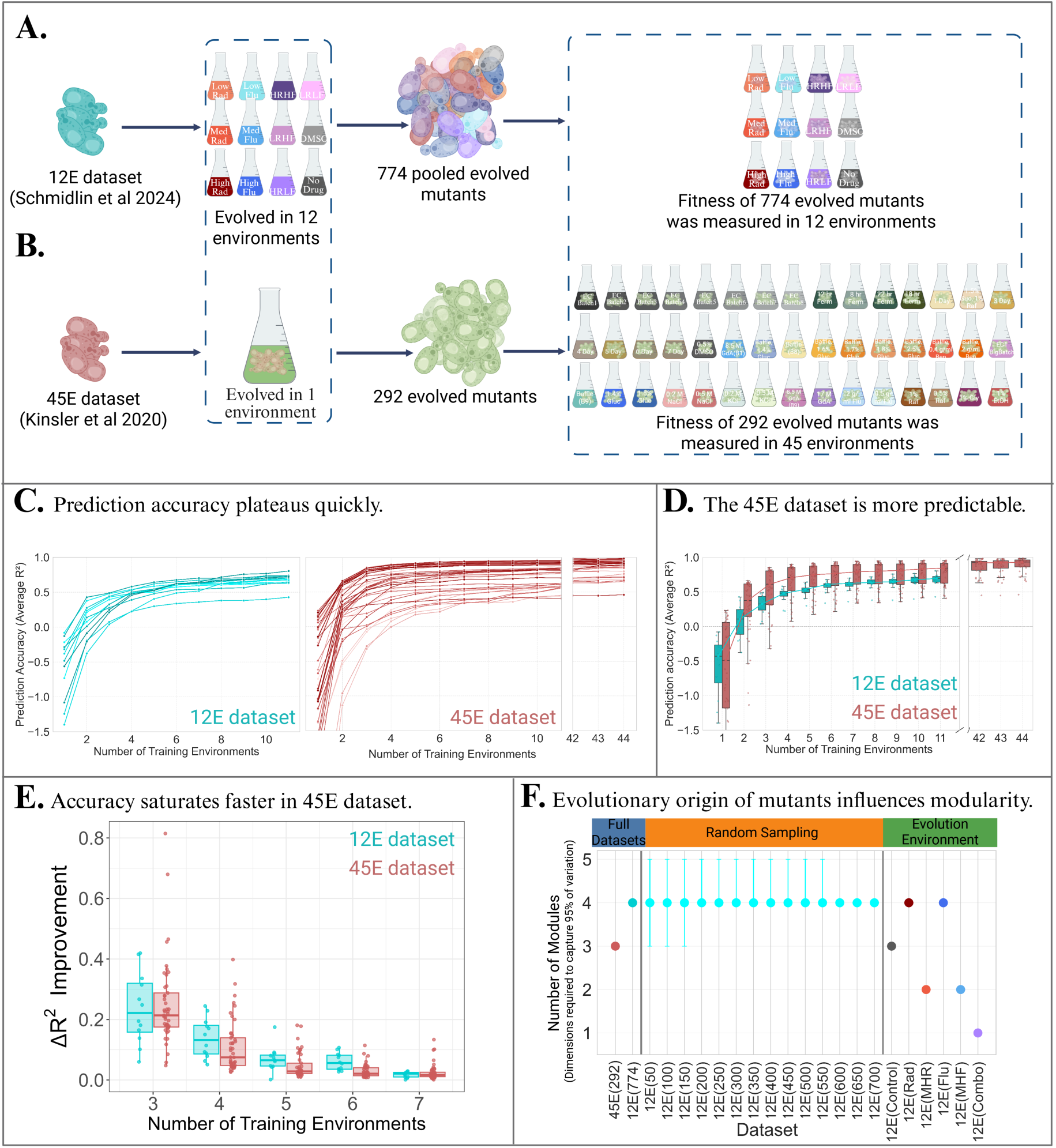
Prediction accuracy saturates rapidly, revealing a modular genotype-phenotype map where dimensionality depends on both mutant and environment diversity. **(A,B)** Schematic comparison of the two fitness datasets analyzed here.**(A)** In the 12-environment (12E) dataset, fitness was measured for 774 mutants across 12 drug environments. These mutants were pooled from 12 different evolution experiments each involving a different drug concentration or combination. **(B)** In the 45-environment (45E) dataset, fitness was measured for 292 mutants across 45 various environments. These mutants originated from a single selection pressure, which was glucose limitation. **(C)** To test how prediction accuracy depends on training data, the number of training environments, *k*, was varied from 1 to *n* — 1, where *n* is the total number of environments. For each value of *k*, 100 iterations were performed in which k environments were randomly selected for training, and predictive accuracy (average R^2^) was evaluated on the held-out environments and averaged across iterations. Training mutants were chosen as in previous work (Kinsler et al. 2020), using a small but balanced subset; this differs from Fig. 2, where training mutants were selected at random. Curves are shown separately for each environment where fitness was surveyed in the 12E dataset (cyan) and the 45E dataset (red). Prediction accuracy increased rapidly upon addition of the first few training environments and then plateaued in both datasets. This rapid saturation indicates that a small subset of environments captures the majority of the information needed to predict fitness. **(D)** Comparison of prediction accuracy across the 12E and 45E datasets. Boxplots summarize the distribution of average R^2^ values across held-out test environments. In general, prediction ac-curacy was higher for the 45E dataset than for the 12E dataset, indicating that fitness in the 45E dataset is generally easier to predict from training data. **(E)** Improvement in prediction accuracy, Δ*R*^2^, obtained by adding each training environment. Shown here are training-set sizes 3–7, where differences between datasets are most apparent. Boxplots summarize the distribution of Δ*R*^2^ values across 100 randomized iterations for the 12E dataset (cyan) and the 45E dataset (red). Improvement declined toward zero in both datasets as more environments were added, but the decline was faster in the 45E dataset, indicating earlier saturation of predictive information. **(F)** Effective modularity of the fitness matrices, measured as the minimum number of SVD dimensions required to explain 95% of the variance. Left panel shows that the full 45E dataset required fewer dimensions (3) than the 12E dataset (which requires 4). Middle panel shows results from 100 repeated random subsamplings of 774 mutants from the 12E dataset at different sample sizes (n = 50–700). Right panel shows (non-random) subsets of 12E mutants selected based on which of the 12 evolution environments they originated from. These results indicate that effective dimensionality is not explained by mutant sample size alone and depends on the diversity and evolutionary origin of the mutants.

This behavior supports a modular view of the genotype–phenotype–fitness map (Kinsler et al. 2020), in which a limited number of inferred phenotypic axes consistently govern fitness across diverse conditions. Once those shared phenotypic modules are learned, adding more environments yields little additional predictive power. At the same time, the plateau exposes a fundamental limit: beyond a small amount of training data, prediction accuracy no longer improves. This ceiling could arise for at least two reasons. First, it may reflect an inherent limitation of linear models such as SVD, which cannot capture nonlinear relationships between phenotypes and fitness, including threshold or saturation effects known to shape adaptive landscapes (Otwinowski et al. 2018; Perfeito et al. 2011). Second, as explained above and emphasized by Ghosh et al. (Ghosh et al. 2025), not all phenotypic functions contribute to fitness in every environment. Thus, some remain hidden from SVD models unless the environments where they matter are represented in training. In this “limiting functions” view, expanding environmental diversity should reveal these latent modules and, in turn, improve prediction accuracy in the contexts where those modules constrain fitness. If that logic holds, then datasets spanning more diverse environments should require more training environments to reach saturation in predictive power.

Yet, when we compared datasets, the opposite occurred. Fitness in the larger, 45-environment dataset is generally easier to predict (Figure 3D). Further, a direct comparison of how prediction accuracy improves with each additional training environment shows that, beyond the third added environment, gains in the 45-environment dataset were consistently smaller than in the 12-environment dataset (Figure 3E). In other words, the 45-environment dataset required fewer training environments to saturate prediction accuracy than did the 12-environment dataset. This pattern is also apparent in figure 3C: prediction accuracy in the 45-environment dataset (red) rises more steeply with the number of training environments and plateaus earlier than in the 12-environment dataset (blue). This contradicts the expectation that broader environmental diversity uncovers more phenotypic modules and thus requires more training environments and phenotypic dimensions to predict fitness.

The most straightforward explanation, that the 45 environments are simply more similar to one another, is not likely given that the 45 environment set spans a wide range of conditions, including antifungal drugs, salts, oxygenation levels, carbon sources, and metabolic states (Kinsler et al. 2020; Li et al. 2018). We posit that the key difference lies not in the environments but in the mutants themselves. Our 12-environment dataset includes 774 mutants evolved under many distinct selective pressures, while the 45-environment dataset includes 292 mutants that all evolved under a single one: glucose limitation (Venkataram et al. 2016; Levy et al. 2015). Because latent phenotypes can only be revealed when they are perturbed, a model trained on a genetically uniform set of mutants will fail to detect modules that those mutations do not affect. If the mutants in the 45-environment dataset collectively perturb fewer phenotypes, then SVD applied to that dataset should uncover fewer phenotypic modules, even though it spans more environments.

Indeed, this is what we observe. When we ask how many phenotypic dimensions are required to capture 95% of fitness variation in the full datasets (with no data held out), that number is smaller (three) for the 45-environment dataset and larger (four) for the 12-environment dataset (Figure 3, S8) When we hold out data and systematically add it back, the number of inferred phenotypic dimensions rises faster in the dataset with more mutants, not the one with more environments (Figure S8). These differences are not due to sample size, down-sampling the 774-mutant dataset does not alter the number of phenotypic dimensions required to capture 95 percent of the variation in fitness (Figure 3F; middle, see also Figure S9). However, these differences are indeed affected by mutant diversity. Down-sampling the 774-mutant dataset to include only mutants that originated from a particular selection pressure often reduces the number of phenotypic dimensions required to capture 95 percent of variation in fitness (Figure 3F; right side, see also Supplementary Figure S10). This highlights that mutant diversity, in addition to environmental diversity, can drive the discovery of new phenotypic dimensions.

This observation reflects something deeper, and wonderfully circular, about evolution. Selection has shaped cellular systems to be modular, and thus the solutions that evolution discovers are themselves modular. Mutants isolated from a single selective pressure tend to perturb a smaller number of phenotypic modules than mutants drawn from twelve distinct selective pressures. Still, in both cases, a small number of shared phenotypic modules underlie fitness. We next sought insights about those phenotypic modules by asking which environments reveal them and which mutants define them.

### 2.3 Interpreting phenotypic modules by examining informative environments and mutants

Having shown, like previous studies (Kinsler et al. 2020; Ghosh et al. 2025; Tenaillon et al. 2012), that the map from genotype to phenotype to fitness appears to be modular, we next ask about the underlying biology that is captured by each phenotypic dimension. To better understand the inferred phenotypic dimensions that drive fitness in multidrug environments, we asked which single-drug environments contribute most to predictive accuracy. Our results demonstrate that individual environments can disproportionately reveal specific phenotypic modules. Specifically, low-dose fluconazole exposes a distinct axis of gene-by-environment interaction that shapes multidrug fitness.

Fluconazole-containing environments were the most informative in general. Removing all fluconazole environments from the training set caused a steep drop in predictive accuracy, while removing radicicol had more modest effects (Figure 4A). The most dramatic impact came from low-dose fluconazole (4 *µ*g/ml). Removing this single-drug environment substantially reduced prediction accuracy in double-drug environments containing low-dose fluconazole; for example, adding it back increased *R*^2^ by ∽0.2 for the LRLF condition (Figure 4B).

**Figure 4:**
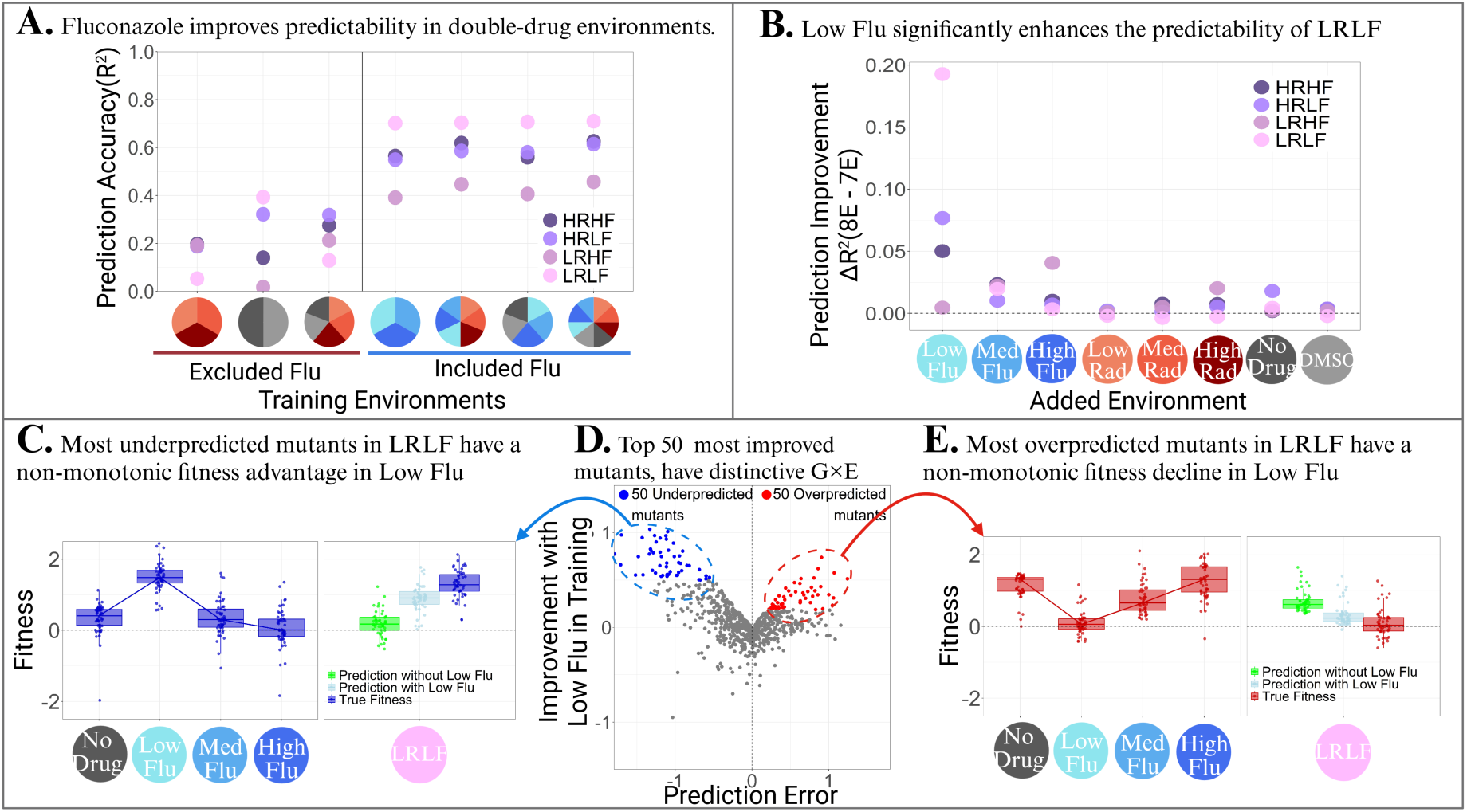
Low-dose fluconazole (4*µ*g/ml) is a uniquely informative environment. To identify which environments provide the most information for predicting fitness in double-drug combinations, we systematically eliminated environments from the training datasets. **(A)** Prediction accuracy improves when fluconazole (Flu) environments are included in the training set. Accuracy was quantified as *R*^2^ between predicted and measured fitness, averaged over 100 training runs, where in each run a random set of 100 mutants was used to predict the fitness of the remaining 674 (as in Figure 2). **(B)** The specific contribution of each single-drug environment to prediction accuracy in each double-drug environment. The y-axis shows the improvement in *R*^2^ when a given environment on the x-axis is added to a base training set of 7 single-drug environments. Adding Low Flu (4 *µ*g/ml) provides the highest improvement in prediction accuracy, specifically for the LRLF test environment (light pink). **(C)** Fitness profiles of the 50 most underpredicted mutants (points) when Low Flu is removed from the training set display a non-monotonic response to fluconazole concentration, with a distinct fitness advantage in Low Flu. The model trained without Low Flu fails to capture this advantage and underpredicts fitness in LRLF (green). Including Low Flu in the training set allows the model to better approximate fitness in LRLF (light blue). **(D)** Relationship between prediction error when Low Flu is excluded from training, and prediction improvement when Low Flu is included (for the LRLF condition). Colored points display the 100 mutants for which predictions improve the most upon adding Low Flu to the training set (blue when excluding Low Flu leads to underprediction, red when it leads to overprediction). **(E)** Fitness profiles of the 50 most overpredicted mutants from panel D. These mutants exhibit an opposite non-monotonic pattern to those in panel C, with a fitness decline in the Low Flu environment compared to other Flu concentrations. The model trained without Low Flu misses this decline and overpredicts their fitness in LRLF (green).

To understand why low fluconazole had such a strong effect, we examined the mutants whose predicted fitness changed the most when low fluconazole was excluded from the training data. Two strikingly different groups emerged (Figure 4C–E). The first group, mutants underpredicted without low fluconazole, had high fitness only at low fluconazole (4 *µ*g/ml) but not in higher doses (Figure 4C, blue). The second, mutants overpredicted without low fluconazole, showed the opposite pattern, performing poorly only at low doses but well in all others (Figure 4E, red).

These mirror-image, nonmonotonic dose–response curves suggest that low-dose fluconazole is not simply a weaker version of drug stress but engages a distinct cellular regime. Similar dose-specific shifts have been observed in molecular studies, where low and high drug concentrations activate qualitatively different physiological responses (Hill et al. 2015; Rosenberg et al. 2018; Berry and Gasch 2008; Pinheiro et al. 2021). Here, we uncovered this unique biology directly from an abstract genotype–fitness map where predictive performance pinpoints low-dose fluconazole as uniquely informative.

Consistent with the idea that low-dose fluconazole triggers a distinct cellular state, mutants with nonmonotonic fluconazole dose–response curves recur across our independent analyses (Figure S11, S12), and prior work has shown that different adaptive lineages emerge at low versus high fluconazole concentrations (Hill et al. 2015; Rosenberg et al. 2018; Harrison et al. 2014). Including low fluconazole consistently improves multidrug prediction across model variants (Figure S13), and fitness in this environment is least correlated with any other condition (Figure S14), underscoring the unique role of low fluconazole in defining a key axis of the genotype–phenotype–fitness map. Together, these findings illustrate how abstract, low-dimensional models can be used not only to predict evolutionary outcomes, but to expose the environments and mutations that define key phenotypic modules.

### 2.4 Our map is sparse, but more asymmetric than other datasets

So far, our analyses reveal that the genotype–phenotype–fitness map is modular in that fitness variation can be captured by a few shared phenotypic axes. We next ask whether these axes are broadly or sparsely connected (Figure 1B). Whereas modularity explains why fitness variation can be captured by a few shared dimensions, sparsity determines how broadly those dimensions are used, whether each mutation or environment engages many modules or just a few. In a sparse system, pleiotropy is limited: each mutation changes only a subset of the phenotypic modules. Further, each phenotypic module matters for fitness in only certain environments. This kind of sparse organization is theorized to arise to offset the “cost of complexity,” the idea that when a mutation affects many traits simultaneously, it can make adaptation slower or less efficient (Orr 2000). At the same time, sparse structure may support evolvability, allowing different parts of an organism to change and adapt more independently (Wagner et al. 2007; Wagner and Altenberg 1996). Thinking about sparsity in this way provides a framework for understanding whether adaptation to the drug environments we study here occurred through a handful of distinct routes or through broad, system-wide changes.

Here we study the degree of sparsity following previous work (Petti et al. 2023). First we applied SVD to the complete matrix of 774 mutant fitness values across 12 environments, without holding out any data (Figure S16). Then we applied a sparsity-constrained variant of SVD, Sparse Structure Discovery (SSD) (Petti et al. 2023) (Figure 5, S16). SSD enforces sparsity by pruning links in the genotype–phenotype–fitness map (Figure 5B; grey arrows). This improves interpretability by highlighting which links are most meaningful.

**Figure 5:**
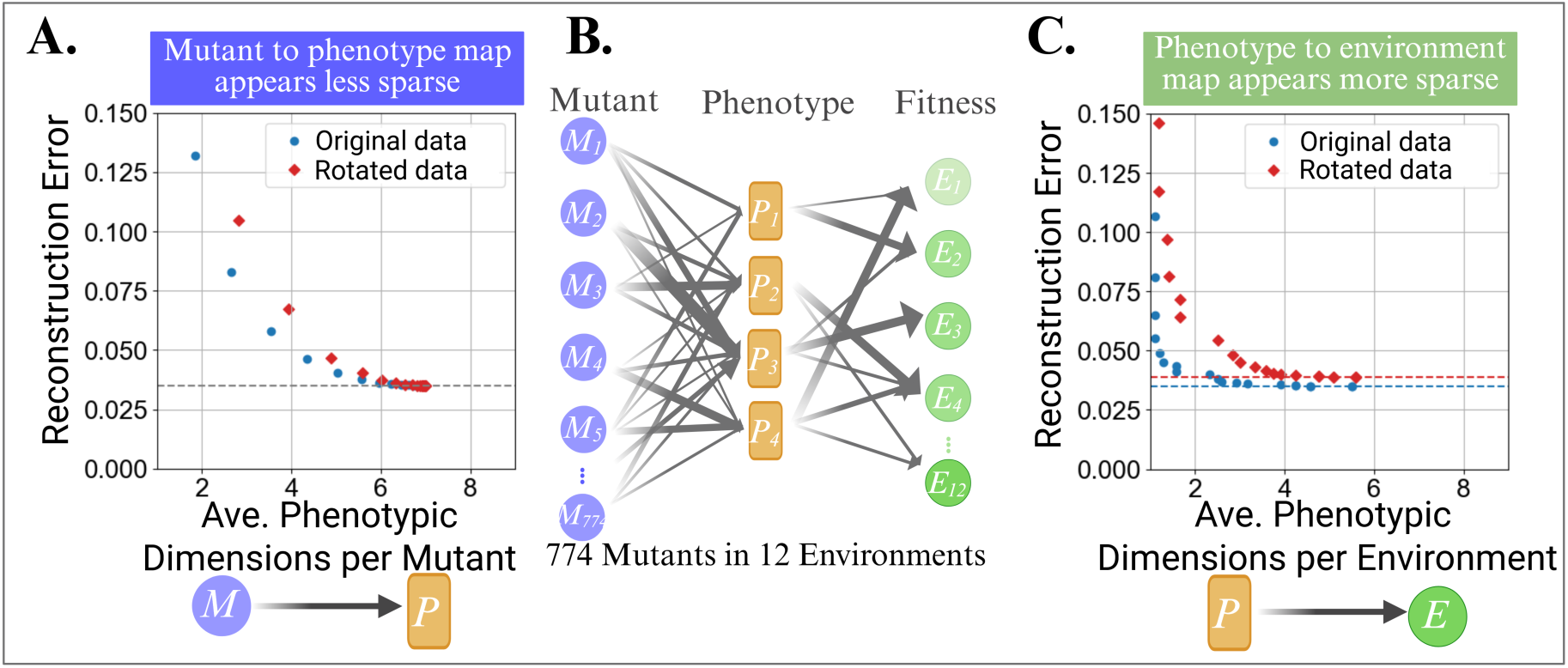
A rotation test supports a less sparse mutant-to-phenotype map but a sparser phenotype-to-environment map. To test for inherent sparsity in our experimental dataset, we applied a rotation test using a sparse discovery algorithm (SSD) (Petti et al. 2023). The rotation test tries to find a sparse, low-error solution for the original fitness matrix, F, versus a rotated matrix (F r = FO), which preserves the correlational structure but removes any special sparse basis. A gap between the solution curves for the original and rotated data indicates that the original data are aligned with a sparse basis. **(A)** The rotation test versus original data for the mutant-to-phenotype mapping. **(B)** A conceptual genotype-phenotype-fitness map that represents the amount of sparsity found within our dataset. Many grey arrows connect mutants to phenotypes, while fewer connect phenotypes to environments, depicting how the latter mapping is sparser. **(C)** The rotation test versus the original data for the phenotype-to-environment mapping. Each point represents a solution for a different regularization parameter. This plot shows the trade-off between reconstruction error and the average number of inferred phenotypes per environment. The original data can be explained by a sparser model (fewer phenotypes per environment) without reducing accuracy, supporting sparse structure on the environment side of the map.

In datasets with truly sparse structure, random rotations of the fitness matrix “F” (Figure 2A) will destroy the alignment between mutants (or environments) and the inferred phenotypic dimensions. As a result, more inferred dimensions are required to achieve similar reconstruction accuracy to the original, unrotated data. This is indeed what we observe. In both the mapping from mutants to phenotypic dimensions (Figure 5A), and the mapping from phenotypic dimensions to environments (Figure 5C), there is a gap between the reconstruction errors on the original versus rotated data. The gap is more apparent in the mapping from phenotypes to environments, suggesting a sparse structure, where not every phenotype affects fitness in every environment. This corresponds to fewer grey arrows connecting phenotypes to environments in our conceptual model (Figure 5B).

Interestingly, rotation tests had less effect on the alignment from mutants to inferred phenotypic dimensions (Figure 5A). This smaller gap between the original and rotated data here suggests that the mapping from mutants to phenotypes is less sparse. In other words, pleiotropy is present, such that each mutation affects multiple phenotypic dimensions, as depicted by the many grey arrows connecting mutants to phenotypes in our conceptual model (Figure 5B). This conclusion is also supported by directly comparing the number of edges per node required to capture the same amount of variation in fitness. Nearly four phenotypic dimensions per mutant are required to capture 95 percent of variation in fitness, while less than two are required per environment (Figure 5). This asymmetric pattern, higher connectivity on the mutation-to-phenotype side, but greater sparsity on the phenotype-to-environment side, departs from most previously studied maps in which both sides showed sparse structure (Petti et al. 2023). Because the mutant and environment sides of the map have very different dimensions (774 mutants vs. 12 environments), we interpret the relative gap sizes cautiously. Synthetic calibration at the same matrix dimensions shows that sparse structure can be recovered under this sampling structure, but also supports using the rotation test primarily as a within-mode diagnostic rather than as a direct numerical ranking of sparsity across modes (Figure S15). One possible explanation for the relatively higher pleiotropy (and lower sparsity) on the mutant-to-phenotype side is that it may reflect the strong selection pressure under which these mutants evolved. Previous work and theory suggest that pleiotropic mutations are more likely to be favored under strong selection pressure (Orr 2000; Fisher 1930), such as antimicrobial drugs used here. More generally, both sides of the map exhibit sparser structure than random rotations, indicating that the inferred structure may reflect a limited set of underlying biological processes.

### 2.5 Phenotypic axes correspond to gene-by-environment interaction patterns

Finally, to connect the modular and sparse structure of the genotype–phenotype–fitness map to underlying biology, we examined which mutants and environments define each phenotypic axis. The phenotypic axes recovered from SSD largely overlapped with those from standard SVD (Figure S16), regardless of which sparse solution we chose. This overlap suggests that the main low-dimensional structure is stable across related decompositions, even though the precise orientation of SVD axes is rotationally ambiguous. This robustness is significant because it suggests the low-dimensional structure we identified is not a fragile statistical abstraction but a primary feature of our studied system.

Unlike other studies (Petti et al. 2023), the phenotypic dimensions we recover are difficult to interpret in terms of discrete cellular functions, perhaps due to low sparsity in the mapping from mutants to inferred phenotypes. For example, although 35 of our 774 mutants perturb regulators of multidrug efflux, there is no single phenotypic dimension that is affected by exclusively these mutants. They predominantly fall into group 2 (Figure 6A; green) and contribute strongly to phenotypic dimension 1 (Figure 6A-B). But consistent with lower sparsity, many other mutants have similar effects on this phenotypic dimension. These effects scale with average fitness advantage across drug environments. Thus, phenotypic dimension 1 seems to capture a common type of gene-by-environment interaction (G×E) whereby fitness is high in environments that contain drugs, and low in environments that do not (Figure 6B).

**Figure 6:**
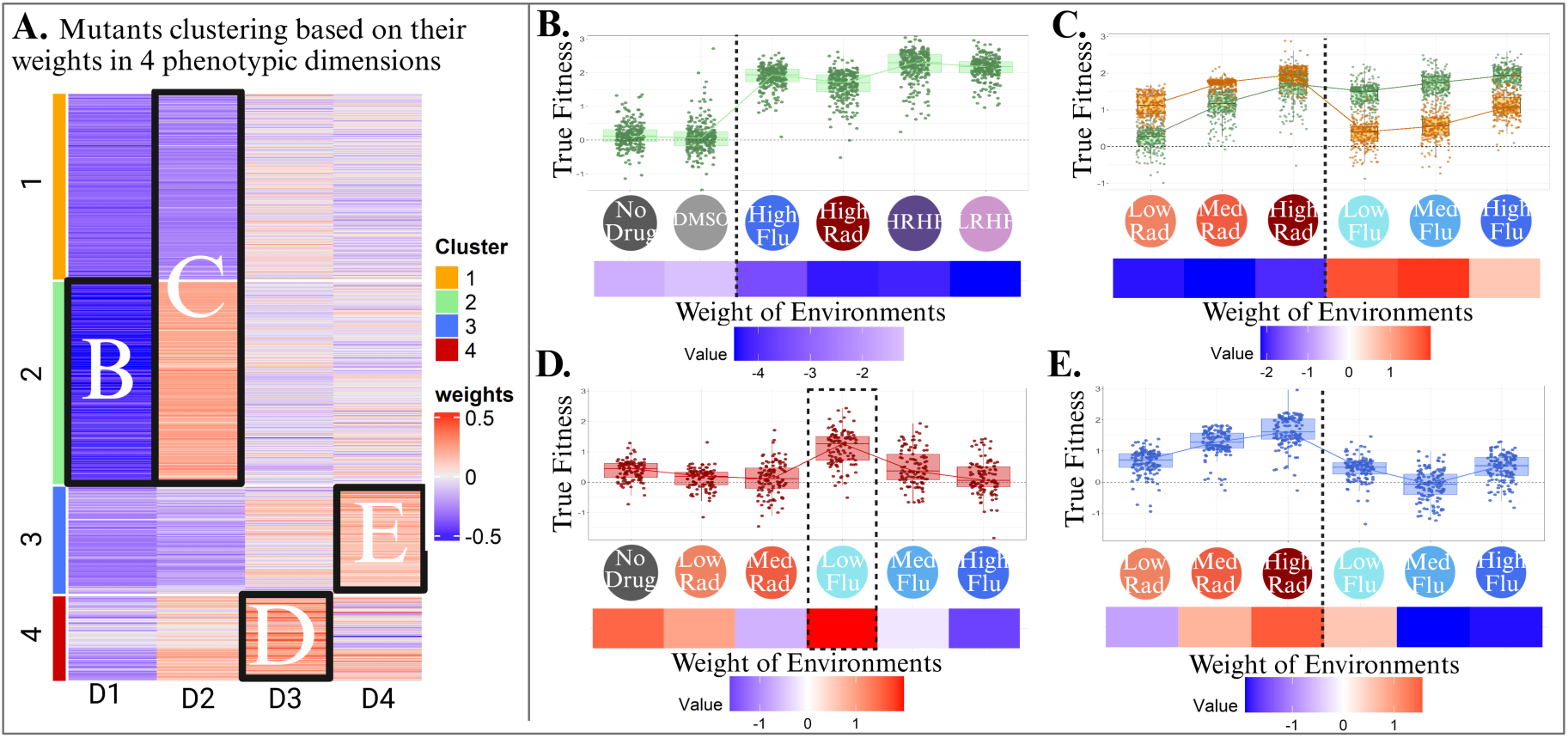
A four-dimensional model decomposes the fitness matrix into phenotypic axes that capture gene-by-environment interactions. Singular Value Decomposition (SVD) was applied to the complete fitness matrix of 774 mutants across all 12 environments to interpret the underlying structure of the genotype-phenotype map. **(A)** A heatmap of the contribution of all 774 mutants to each inferred phenotypic dimension (red vs. blue indicate strong contribution in opposite directions). Mutants are clustered based on their weight profiles (calculated based on 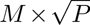). Boxes highlight the mutant clusters detailed in panels B–E. **(B)** This phenotypic dimension is most strongly affected by mutants that have fitness advantages across environments containing drugs that are lost in environments without drugs. Mutants with amino acid changes in the PDR1 and PDR3 genes, which are regulators of drug efflux pumps, largely fall into this category. **(C)** This dimension is strongly influenced in opposite directions by two groups of mutants: one that resists flu more than rad (green), and another that resists rad more than flu (orange). **(D)** This dimension captures a specific response to low-dose fluconazole, which has the strongest positive environmental weight. Mutants with high weights in this dimension show a distinct fitness advantage in this specific environment. **(E)** This dimension reflects a distinct nonmonotonic response to fluconazole: fitness is lowest at intermediate concentrations. This pattern differs from the nonmonotonic responses in Figure 4, where mutants are instead most sensitive in low fluconazole.

Similarly, the other phenotypic dimensions seem to map to G×E interactions more so than underlying physiological mechanisms by which mutants affect fitness. For example, phenotypic dimension 2 separates mutants based on relative advantage in fluconazole versus radicicol (Figure 6C). Dimensions 3 and 4 capture patterns of G×E related to changes in fluconazole concentration (Figure 6D–E).

These results suggest that the extracted dimensions likely reflect shared fitness outcomes of mutations, rather than the unique cellular process that underlie those outcomes. Interpreting the underlying biological mechanisms that link genotype to fitness may therefore require integrating additional information, such as mutant genotypes or transcriptional profiles. Still, the fact that just four dimensions explain the vast majority of fitness variation, despite the genetic diversity present across this collection of 774 mutants, implies that genotype-by-environment interactions are canalized. In other words, every genetic perturbation does not have a distinct pattern of G×E. Instead, a small number of G×E patterns appear to emerge that describe most mutants (Schmidlin et al. 2024). This canalization might arise from constraints imposed by regulatory network architecture or trade-offs in resource allocation, which may in turn restrict the evolutionary routes available for drug resistance. Whatever the cause, this structure is encouraging because it shows that even though mutations are inherently pleiotropic, in the sense that they can influence countless molecular-level traits, their fitness effects often funnel through a small number of shared axes. This implies that mapping the underlying molecular mechanisms onto these axes, identifying the core processes that shape fitness, and ultimately predicting evolution, may be more tractable than one might expect given the apparent complexity of all that a mutation could, in principle, alter inside a cell.

## 3 Discussion

Our results show that, despite extensive genetic and environmental complexity, the fitness effects of hundreds of antifungal-resistance mutations can be captured by a strikingly low-dimensional structure. A small number of inferred phenotypic axes explain most variation in mutant fitness across drugs and doses, and axes learned from single-drug environments generalize to predict fitness in drug combinations. This supports a strongly modular genotype–phenotype–fitness map. Rather than each mutant having its own idiosyncratic genotype-by-environment (G×E) pattern, evolution in multidrug environments is governed by a few repeatable ways that mutants interact with drugs.

Our results echo ideas from the omnigenic perspective on complex traits, which holds that mutations in many genes can shape trait variation through dense regulatory networks (Boyle et al. 2017; Liu et al. 2019). In our case, mutational effects did not scatter unpredictably but instead collapsed onto a small set of shared axes. A recent quantitative omnigenic model formalized this principle, showing that network-level complexity can be summarized in low-dimensional, interpretable structures (Ružičková et al. 2024). The key difference is that while the quantitative omnigenic framework is a bottom up approach deriving axes from regulatory network topology, our top-down SVD approach infers them from observed fitness patterns alone. Yet both converge on the idea that molecular complexity funnels through a limited set of modules that enable prediction.

In our dataset, the architecture of this modular map is asymmetric. Our sparsity analysis shows that the mapping from mutations to phenotypic modules is less sparse, consistent with higher pleiotropy, while the mapping from phenotypes to environments is more sparse. To understand why, we step back to foundational theory. Under Fisher’s Geometric Model, adaptation to novel environments is predicted to proceed via large-effect, often pleiotropic mutations, whereas stabilizing selection should favour smaller changes (Fisher 1930; Orr 2000; Wagner and Zhang 2011; Welch and Waxman 2003). Empirical evolution studies echo this pattern: early adaptation to novel or harsh environments often proceeds via large-effect, highly pleiotropic mutations in global regulators or metabolic hubs (Saxer et al. 2014; Conrad et al. 2011; McCloskey et al. 2018), whereas subsequent adaptive steps and adaptation under more benign environmental shifts tends toward smaller-effect, more modular changes (Kinsler et al. 2024; Bomblies and Peichel 2022). This framework suggests that the amount of pleiotropy (and thus sparsity) on the mutant-to-phenotype side of the map depends on the selection regime under which the mutations evolved. Denser mutant-to-phenotype mappings may be common under adaptation to novel environments or strong stressors. Indeed, a denser mutant-to-phenotype map was also observed in another study of adaptation to a novel environment, but only after eliminating a class of mutations that is not present in our dataset (see Figure S4 in Petti et al. 2023 (Petti et al. 2023)). In contrast, the consistent sparsity of phenotype-to-environment mappings across ours and others datasets (Petti et al. 2023) may reflect long-term selection for mutations with modular effects on phenotype and fitness. In this way, our findings not only support Fisher’s geometric model–derived expectations about pleiotropy and effect size, but also suggest that the architecture of genotype–phenotype–environment maps is shaped by both past selective histories and present-day selective contexts.

Our analysis also highlights how the inferred dimensionality of any given genotype-phenotype map depends as much on which mutants are assayed as on which environments are sampled. We showed that increasing genetic diversity can reveal new phenotypic modules even when environmental coverage is modest, whereas expanding environmental panels without diversifying mutants may uncover relatively fewer modules. This has practical implications for experimental design: to map genotype–phenotype–fitness structure efficiently, it may be important to study many kinds of mutants as well as many environments. Even with pervasive pleiotropy, mutations differ in how strongly they affect each phenotypic module, so sampling diverse mutants is essential for resolving the underlying structure.

Low dimensional models can do more than reveal the structure of the genotype-phenotype relationships, here we show that they can also be used as discovery tools. Low-dose fluconazole emerged as a particularly informative condition that likely perturbs a unique physiological response. Excluding it from the training set degraded predictive performance despite other fluconazole concentrations being included in the model. This mirrors molecular evidence that low and high drug concentrations can induce distinct physiological states, rather than a simple graded stress response. Crucially, we detected this potentially unique physiological response, and the specific mutants that define it, purely from patterns of predictive success and failure.

These findings have implications for the management of antimicrobial resistance, particularly for collateral sensitivity cycling. Our data show that tradeoffs are not uniform; mutants selected by the same drug can reside in opposing regions of the fitness landscape in a second environment. Consequently, a ‘one-size-fits-all’ second-line drug may suppress one cluster of resistant mutants while favoring another. Effective multidrug therapies may therefore need to move beyond population averages and towards ‘cluster-specific’ strategies, where the specific phenotypic fingerprint of the resistant sub-population informs the next move in the therapeutic sequence.

Looking forward, while our linear model successfully captures the bulk of the fitness variation, its specific failures are as illuminating as its successes. The identification of environments and mutant subsets that defy prediction points to potential non-linear interactions that shape fitness. Several questions remain. How general is the low-dimensional modular structure across timescales, selective pressures, or organisms (Geiler-Samerotte et al. 2020; Kinsler et al. 2020; Wagner et al. 2007; Orr 2000)? What molecular or physiological variables underlie the axes we infer, and how can we map them to interpretable traits (Manrubia et al. 2021; Petti et al. 2023; Martí-Góomez et al. 2025)? To what extent do nonlinear or environment-specific modules emerge in more complex conditions (Ghosh et al. 2025; Avasthi et al. 2023)? Addressing these questions will require integrating multi-omics data and nonlinear modeling frameworks. Such approaches may bridge the gap between predictive low-dimensional structure and the underlying mechanisms, illuminating how mutations ripple through networks to produce fitness effects.

## 4 Methods

### 4.1 Fitness data, yeast strains and experimental design

The yeast strains and experimental methods employed in this study were previously described in detail (Schmidlin et al. 2024). Briefly, all yeast strains originated from a single ancestral “landing pad strain” (SHA185), a MAT ↵ haploid strain characterized by the genetic background: *ura3*Δ*0, ybr209w::Gal-Cre-KanMX-1/2URA3-loxP-Barcode-1/2URA3-HygMX-lox66/71*.

To generate the mutants, around 300,000 unique neutral DNA barcodes were introduced into this strain using a Cre-lox recombination system (Levy et al. 2015). These barcoded yeast were evolved in 12 environments. These 12 evolution experiments were conducted in a glucose-limited M3 medium lacking uracil, supplemented with fluconazole, radicicol, or DMSO at the following concentrations: Low Flu (4 *µ*g/ml), Med Flu (6 *µ*g/ml), High Flu (8 *µ*g/ml), Low Rad (5 *µ*M), Med Rad (15 *µ*M), High Rad (20 *µ*M), LRLF (5 *µ*M + 4 *µ*g/ml), LRHF (5 *µ*M + 8 *µ*g/ml), HRLF (15 *µ*M + 4 *µ*g/ml), HRHF (15 *µ*M + 8 *µ*g/ml), 0.5% DMSO (0.5%) and No Drug (no supplement). Drug concentrations were chosen to apply mild selective pressures without dramatically reducing barcode diversity (Schmidlin et al. 2024). Herein, we refer to the 4 environments containing both fluconazole and radicicol as *“double-drug environments”* and all others as *“single-drug environments.”* In previous work, adaptive yeast mutants, isolated at intermediate evolutionary time points to ensure maximal barcode diversity, were combined from all 12 evolution experiments into a single pooled population. Then they were competed against an ancestral strain in each of the 12 aforementioned experimental conditions. Since two replicate experiments were performed in each condition, 24 total fitness measurement experiments were included. Fitness in each experiment was inferred from log-linear changes in each mutant’s barcode frequency, over time, relative to the ancestor. Barcode frequencies were measured by Illumina sequencing after sequential batch culture transfers. Only barcodes meeting strict sequencing coverage criteria (≥500 counts across time points) were included. Fitness measurements were averaged for each barcode across replicates, resulting in a dataset comprising fitness values for 774 adaptive yeast mutants across all 12 conditions.

Sixty of these 774 adaptive mutant lineages have been previously sequenced (Schmidlin et al. 2024). Most differ from the ancestor by only one or two amino acid changes. Thirty five have (different) amino acid changing mutations to either PDR1 and PDR3, known regulators of a drug efflux pump (PDR5).

These 774 lineages have also been previously clustered into distinct classes based on their G×E interactions using a Gaussian Mixture Model and UMAP (Schmidlin et al. 2024). For example, lineages with mutations in PDR genes tend to cluster into a class with high fitness in most drug environments, but reduced fitness in drug-free environments (Schmidlin et al. 2024). Whereas, another class of lineages exhibits high fitness specifically at low fluconazole concentrations, but low fitness at higher doses. We used these previously defined G×E-based clusters to construct balanced training sets of mutants. Specifically, when selecting mutants for analyses requiring a fixed training set (e.g., Figure 3), we sampled mutants to preserve the proportion of each cluster in the data, thereby capturing diverse and representative patterns of environmental response.

For the comparison dataset with 45 environments, a balanced mutant set had already been defined (Kinsler et al. 2020). In that case, balance was achieved by selecting mutants across distinct mutated genes rather than across G×E interaction classes. This comparison dataset (Kinsler et al. 2020) was collected following a very similar experimental design to the focal dataset (Schmidlin et al. 2024) in terms of the barcoded library construction, the number of generations of evolution, the number of differences between each barcoded strain and the ancestor, and the way fitness was measured in each environment.

For a complete description of yeast strain construction, barcode library generation, and detailed fitness assay protocols, please refer to Kinsler et al. 2020 (Kinsler et al. 2020) and Schmidlin et al. 2024 (Schmidlin et al. 2024).

### 4.2 Dimensional reduction of the fitness matrix via singular value decomposition (SVD)

We applied singular value decomposition (SVD) to reduce the dimensionality of a fitness matrix consisting of 774 mutants measured across 12 environments. In matrix notation,

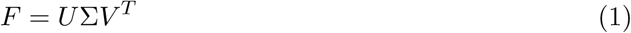

where *F* is the *N* × *E* fitness matrix (*N* = 774, *E* = 12), *U* is an *N* × r matrix of left singular vectors (mutant loadings), *V* is an *E* × *r* matrix of right singular vectors (environment loadings), and Σ is an *r* × *r* diagonal matrix of singular values.

In this framework, the columns of U describe how individual mutants load onto each latent dimension, while the columns of V describe how environments load onto those same dimensions. The diagonal entries of Σ quantify the amount of variation explained by each dimension. Together, these matrices define abstract phenotypic dimensions that summarize systematic patterns of genotype–environment interactions.

Depending on the analysis, different combinations of mutants and environments were used to train the SVD model. These differences are described in the main text, the figure legends, and below. All analyses were conducted using custom Python scripts. The full code, including a Jupyter notebook demonstrating the SVD workflow, has been made publicly available at https://github.com/Russellrezagh/SVD-SSD-genotype-phenotype-mapping.

### 4.3 Bi-cross-validation of SVD models

In most figures, a bi-cross-validation approach was used to assess the predictive performance of the SVD-based model (Owen and Perry 2009; Kinsler et al. 2020). This approach involves choosing a subset of mutants and environments as the training data, building a low dimensional model, and then using that model to predict the fitness of the held out mutants and environments, which are referred to as the test data.

In bi-cross-validation, the fitness data matrix (matrix F) is divided into four submatrices: (submatrix A) test mutants in test environments, (submatrix B) test mutants in training environments, (submatrix C) training mutants in test environments, and (submatrix D) training mutants in training environments. SVD is applied to the training submatrix (submatrix D), producing singular values and singular vectors that represent latent dimensions in the training data. Predictions for the held-out submatrix (submatrix A) are then generated using the relationship

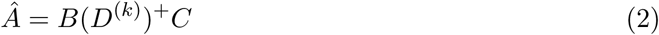

where *D*^(*k*)^ is the rank-k approximation of *D*, obtained by retaining the top k singular values and corresponding vectors, and (*D*^(*k*)^)^+^ is its Moore–Penrose pseudo-inverse. This pseudo-inverse step is equivalent to solving a least-squares regression problem, in which the test mutants (matrix B) and test environments (matrix C) are projected into the training-dimensional space identified by SVD.

### 4.4 Predicting fitness in double-drug conditions from single-drug data (Figure 2)

#### Training data

In figure 2, we implemented bi-cross validation (described above) while using eight single-drug environments in the training data. These environments included three concentrations of fluconazole (Low, Medium, and High) and three concentrations of radicicol (Low, Medium, and High), together with two control environments (No Drug and DMSO). We also included 100 mutants in the training data, selected randomly from the full set of 774. The selection of training mutants was resampled 100 times, and all subsequent procedures were repeated each time, to ensure that results did not depend on the particular mutants chosen.

#### Test data

In each iteration, the remaining 674 mutants were held out as the test set, as were the four multidrug environmental conditions (LRLF, LRHF, HRLF, HRHF).

#### Choosing the optimal number of phenotypic dimensions

In each iteration, we chose the number of latent phenotypic dimensions (r) that minimized the mean squared error on the fitness predictions. We chose this approach because prediction accuracy initially improves as dimensions are added because biologically meaningful patterns are captured. However, the prediction accuracy eventually begins to decline with r when additional phenotypic dimensions incorporate noise rather than signal. Therefore, the optimal number of dimensions was identified by minimizing mean squared error (MSE), as in previous work (Kinsler et al. 2020). Because this rank is chosen from the same held-out predictions used to summarize performance, these values should be interpreted as best-rank held-out performance for this matrix rather than as a fully prospective rank-selection procedure. The optimal number of dimensions varied across our 100 iterations (Figure S1), as did the fitness predictions for each held out mutant in each held out environment. We averaged the fitness predictions pertaining to each mutant across all iterations where it was held out. Comparing these averages to the measured fitness values provides the robust estimates of predictive performance that we report in Figures 2D–E.

#### Calculating prediction accuracy

Prediction accuracy is calculated as *R*^2^ of the predicted versus true fitness. *R*^2^ was computed using sklearn.metrics.r2 score from the scikit-learn library in Python. As described in the package documentation, “Best possible score is 1.0 and it can be negative (because the model can be arbitrarily worse). In the general case when the true y is non-constant, a constant model that always predicts the average y disregarding the input features would get a score of 0.0.” Mathematically, the R-squared is defined as

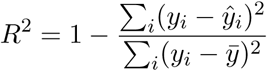

where *y_i_* is the observed value, *ŷ_i_* is the predicted value, and *ȳ* is the mean of the observed values. The term (*y_i_* — *ŷ_i_*) represents the prediction error_P_, that is, the difference between the observed and predicted value for each data point, and Σ*_i_*(*y_i_* − *ŷ_i_*)^2^ is the residual sum of squares. The term (*y_i_* — *ȳ*) represents the deviation of each observed value from the mean of the observed values, and Σ*_i_*(*y_i_* − *ȳ_i_*)^2^ is the total sum of squares or total variance, which measures the total variability of the observed data around their mean. Under this definition, *R*^2^ becomes negative when the residual sum of squares is greater than the total sum of squares, meaning that the model performs worse than the mean of the observed response. Therefore, a negative *R*^2^ indicates poor predictive performance relative to this mean-based baseline.

### 4.5 Evaluating the effect of environmental complexity on predictive power (Figure 3)

To test whether genotype–fitness relationships are organized by a small set of phenotypic modules, we asked how quickly prediction accuracy improves, and then plateaus, as (i) we add more training environments and (ii) more latent dimensions are used in the model. Rapid saturation along either axis is expected if a limited set of phenotypic processes consistently shapes fitness across conditions.

We performed the analysis on two independent datasets including the one that is the focus of the rest of the study (774 drug-resistant mutants × 12 drug environments) (Schmidlin et al. 2024) and one which surveys fitness across a larger number of environments (292 low-glucose-evolved mutants × 45 diverse environments) (Kinsler et al. 2020).

Since we wanted to focus on the effects of altering the number of training environments, we held the number of training mutants constant within each dataset in these analyses. We also balanced the training mutants to contain an even number from different mutants classes, as in previous work (Kinsler et al. 2020). Using balanced mutants reduces the chance that missing a mutant class masquerades as a need for more environments. From the 774 drug-resistant mutants, we chose 100 mutants that evenly represent previously defined mutant clusters with characteristic patterns of G×E (Schmidlin et al. 2024). From the 292 low-glucose-evolved mutants, we chose the same balanced set as was used previously, which includes 60 mutants balanced across genotypes (Kinsler et al. 2020). These training mutant sets were fixed (unlike in Figure 2, where training mutants were repeatedly re-sampled at random).

Here, the training environments were repeatedly re-sampled at random. For each dataset with n total environments, we varied the number of training environments (*k*) from 1 to (*n* — 1). For each *k*, we drew 100 random subsets of *k* environments for training; the remaining *n* — *k* environments were held out for testing. For each iteration, we performed bi-cross validation (see above) choosing the number of latent phenotypic dimensions (r) that minimized the mean squared error on the fitness predictions. We then recorded the corresponding *R*^2^ between predicted and measured fitness for all test mutants in the test environments. Averaging *R*^2^ across 100 iterations for each level of training environments (for *k* = 1 through *k* = *n* 1) gives *R*^2^ for each level of training environments **(Figure 3).** Averaging across all iterations for each number of optimal dimensions (for r = 1 through r = maximum r) gives *R*^2^ for each number of optimal dimensions, which is displayed in panels E and F of Figure S7. To capture both effects together, we also plotted the average *R*^2^ as a function of both *k* and r; these plots show how predictive accuracy rises quickly with additional environments and dimensions before reaching a broad plateau(Figure S7 C,D).

### 4.6 Evaluating the effect of environmental complexity on number of dimensions (Figure S8)

To continue examining the effect of adding training environments on model complexity, we performed a similar analysis to that in Figure 3 in Figure S8. Here, instead of evaluating how predictive power (*R*^2^) changes as we add training environments, we evaluate how the number of dimensions required for a reconstruction of the training data changes as we add training environments (Figure S8). For each number of environments (k = 2 through *k* = *n* — 1), we determined the minimum number of phenotypic dimensions necessary to capture 95% of the data variability. For robustness, each incremental increase in environment count was iterated 100 times, with a different random subset of environments selected for each iteration. Since this was a reconstruction, no mutant sampling was performed, all 774 or 292 mutants were included, depending on the dataset. The number of dimensions required to reconstruct the training data increased with the number of environments more quickly in the 12E dataset (i.e., the dataset pertaining to the 774 drug-resistant mutants × 12 drug environments) (Schmidlin et al. 2024).

The same result was true when we hold the number of environments constant (at n) and test the effect of increasing the number of mutants on model complexity. The 12E dataset pertaining to drug resistant mutants requires more dimensions to capture 95% of the variation in fitness data than does the dataset pertaining to mutants evolved under glucose limitation (Figure S8).

To test if this higher dimensionality was a result of the mixed evolutionary backgrounds in the 12E dataset, we compared the optimal dimensionality of random 12E subsamples against subsets grouped by their evolutionary origin (e.g., those evolved in High Flu vs. Low Rad). While random subsamples (n = 50–700) consistently required 4 dimensions on average to explain 95% of the variance, subsets grouped by evolutionary background yielded distinct optimal dimensionalities (Figure 3, panel F). This indicates that the latent dimensionality of fitness data could be influenced by the specific evolutionary history and diversity of the lineages being studied.

To determine where each mutant evolved, we utilized the metadata from (Schmidlin et al. 2024). Because barcodes were inserted prior to the evolution experiments, some barcodes that mark adaptive lineages were present in multiple evolution environments. Previous work assigned an origin probability to each mutant based on its relative frequency across all evolution experiments where it was detected. To ensure high-confidence mapping, we subsampled this dataset to include only lineages with a >70% probability of belonging to a specific evolutionary background. Using this threshold, we narrowed the 774 lineages down to 563 high-confidence lineages assigned to dominant origins. The number of barcoded lineages assigned to each category is as follows: Control: 123, All radicicol concentrations: 228, MHR (a subset of the former with only medium and high radicicol): 134, All fluconazole concentrations: 142, MHF (a subset of the former with only medium and high fluconazole): 94, Multi-drug combinations, 70.

To further investigate whether the observed complexity in the 12E dataset was driven by the diversity of evolutionary backgrounds, we performed bi-cross validation on the 123 mutants that evolved exclusively in control (DMSO and drug-free) environments. These environments most closely match the single evolution environment from the 45E dataset. We found that when the evolutionary background is restricted to the control conditions, the prediction accuracy pattern (Δ*R*^2^) became more similar to the 45E dataset (Figure S10).

### 4.7 Evaluating each training environment’s influence on prediction accuracy (Figure 4)

To assess how certain types of training environments impact predictive performance of SVD, we trained our model with smaller training sets including some that omitted radicicol, some that omitted fluconazole, some that omitted the control conditions, and some other combinations (Figure 4A). To assess how specific training environments impact the predictive performance of our SVD model, we systematically included and excluded each single-drug environment from the training set and calculated the prediction accuracy using the coefficient of determination (*R*^2^). We assessed the improvement (Δ*R*^2^) in prediction accuracy for each double-drug environment that resulted upon adding individual single-drug environments to the 7-environment training set. We do so 100 times, each time using 100 random mutants in our training set. For each mutant, we averaged the resulting fitness predictions, then calculated the Δ*R*^2^ values between each mutant’s average prediction and measured fitness. The values on the vertical axis of figure 4B represent these averages.

After identifying that including the low fluconazole environment in the training set notably improved predictions in the LRLF condition, we further analyzed which mutants contributed most significantly to this improvement. Specifically, we a identified two groups of mutants: the 50 mutants whose fitness was underpredicted when low fluconazole was excluded and the 50 mutants whose fitness was overpredicted when low fluconazole was excluded. We examined their gene-by-environment interaction profiles and observed characteristic behaviors that echo those detected in an earlier study using different methods (Figure S11) (Schmidlin et al. 2024).

To robustly evaluate the effect of low fluconazole, we extended our drop-out analysis beyond single environments. In addition to testing the effect of adding one environment at a time to the baseline training set, we also systematically added all unique pairs and triplets of environments. For each case, the improvement in prediction accuracy (Δ*R*^2^) was recalculated relative to the 5- or 6- environment baseline (Figure S13).

### 4.8 Using SVD as an interpretation tool rather than for making predictions (Figure 5–6)

We conducted a comprehensive dimensionality analysis using the entire dataset to evaluate the interpretative power of singular value decomposition (SVD) in capturing phenotypic drivers of fitness. Rather than using bi-cross-validation, here we applied SVD directly to the entire fitness dataset, without separating training and test subsets, to identify abstract phenotypic dimensions. We performed SVD on the complete data matrix (774 mutants by 12 environments) and identified the number of phenotypic dimensions needed to explain at least 95% of the observed variation in fitness. Our analysis revealed that just four dimensions were sufficient to capture this threshold of variation. To quantify the overall reconstruction accuracy of our SVD model, we calculated the coefficient of determination (*R*^2^) by comparing the predicted (reconstructed) fitness values against the observed values across the entire dataset. This resulted in an accurate reconstruction (*R*^2^ overall was 0.87) (Figure S16). This reconstruction *R*^2^ is not identical to the fraction of variance explained by the retained singular values, and its value likely reflects both measurement noise and model mismatch. We also computed *R*^2^ values for each individual environment independently to assess how effective our linear dimensionality reduction method works at capturing patterns of fitness variation within each environment (Figure S16). Having reconstructed a comprehensive genotype-phenotype-map that was modular (only four dimensions) we next applied SSD to test it for sparsity.

### 4.9 Analysing sparsity with SSD models (Figure 5)

To enhance our ability to interpret the biological meaning of the inferred phenotypic dimensions, and assess the robustness of the identified dimensions, we sought to resolve the inherent ambiguity in standard SVD. While SVD excels at identifying the optimal low-dimensional subspace that captures the maximal variance in fitness, its solution is rotationally invariant. This means that any orthogonal rotation of its basis vectors (the phenotypic axes) within that subspace is equally valid from a reconstruction standpoint, leaving the specific axes returned by the algorithm biologically arbitrary. To arrive at a more biologically meaningful set of axes, one could introduce a guiding prior that selects for a particular kind of structure.

Here, we adopt the principle of sparsity as a biologically motivated prior, operationalized through the Sparse Structure Discovery (SSD) framework (Petti et al. 2023). This approach assumes that the genotype-phenotype-fitness map is not a dense, all-to-all network. Instead, it posits that fitness in any given environment is shaped by only a small subset of core phenotypic axes (environment-sparsity), and conversely, that any given mutation perturbs only a few of these axes (mutant-sparsity). This framework translates the search for a sparse decomposition into a concrete optimization problem. We seek to find matrices for environmental weights (W) and mutant weights (M) that minimize the objective function:

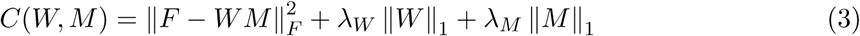

where the first term is the squared Frobenius norm representing the reconstruction error, and the subsequent terms are *L*_1_-norm penalties that enforce sparsity on *W* and *M*, respectively. In the SSD notation used here, *F* is an *E* × *L* matrix, *W* is an *E* × *k* matrix of environment-side weights on phenotypic axes, and M is a *k* × *L* matrix of mutant-side weights on those same axes. For the empirical SSD fits, *k* = 4, matching the four SVD dimensions that captured 95% of the variation. *W* is the SSD analogue of the environment-side loading matrix E in Figure 2A. The regularization parameters, λ*_W_* and λ*_M_*, allow us to tune the trade-off between reconstruction accuracy and the desired level of sparsity.

Crucially, the appropriateness of imposing such a sparsity prior is not an axiom but an empirically testable hypothesis. We evaluated this assumption for our dataset using the rotation test (Petti et al. 2023). This procedure tests whether sparsity is tied to the observed mutant or environment axes rather than being obtainable after an arbitrary rotation. By applying a random orthogonal rotation to the fitness matrix (e.g., *F′ = FO*), we preserve its underlying correlational geometry and SVD reconstruction error while disrupting any privileged sparse basis. If the original data matrix *F* can be approximated by a substantially sparser model than its rotated counterpart *F′* for a given error budget, it provides evidence for an inherent sparse structure.

To perform the rotation test, we represent the data as a fitness matrix *F* ∈ R*^ExL^*, with environments as rows and mutants as columns. This is the transpose of the SVD orientation used above, but it makes the SSD factors align directly with environment weights (W) and mutant weights (M). We then compare SSD solutions obtained from the original matrix *F* to solutions obtained from an orthogonally rotated matrix *F′*. For the mutant-to-phenotype test (Fig. 5A), we rotate the mutant axis by right-multiplying *F* by a random orthogonal matrix, such that *F ^r^* = FO*_L_*. For the phenotype-to-environment test (Fig. 5C), we rotate the environment axis by left-multiplying *F* by a random orthogonal matrix, such that *F′* = *O_E_F*. These rotations preserve the overall second-order structure while removing any privileged sparse basis on the rotated mode.

In the rotation tests, we did not use a single (λ*_W_*, λ*_M_*) pair; rather, each curve in Fig. 5 is a sweep over one regularization parameter while holding the other fixed. In our notation, λ*_W_* penalizes *W* and λ*_M_* penalizes M. For the mutant-to-phenotype rotation test in Fig. 5A, we fixed λ*_W_* = 10*^—^*^4^ and swept λ*_M_* over 25 logarithmically spaced values from 10*^—^*^3^ to 1.5. For the phenotype-to-environment rotation test in Fig. 5C, we fixed λ*_M_* = 10*^—^*^3^ and swept λ*_W_* over 25 logarithmically spaced values from 10*^—^*^4^ to 10*^—^*^2^. These are the same regularization settings used in the previous study of sparsity in an evolution experiment (Petti et al. 2023). Here we applied the same regularization parameter grid to both the original and rotated matrices, ensuring a fair comparison.

Our analysis revealed asymmetry in the structure of our system. The rotation test for the environment-phenotype map demonstrated a clear sparsity-error gap, supporting the interpretation that fitness in each environment is governed by a small number of phenotypic axes (Figure 5C). In contrast, the test for the mutant-phenotype map showed a less striking gap; the original and rotated data were less distinguishable, indicating a less sparse mapping (Figure 5A). This, and subsequent analyses, suggests that the drug-resistant mutations in our dataset have pervasive pleiotropy, with many mutations influencing several of the underlying phenotypic axes. Informed by these results, we leveraged SSD to derive a model that respects this observed asymmetrical structure, yielding a more constrained representation of the genotype-phenotype-fitness architecture (Figure S16).

We also used synthetic calibration to ask how much of the apparent asymmetry could arise from the unequal dimensions of our matrix, specifically having 774 mutants but only 12 environments. Synthetic fitness matrices were generated with E = 12 environments, L = 774 mutants, and *k* = 7 latent phenotypic dimensions using the SSD synthetic generator with Gaussian noise parameter σ = 0.3 (Petti et al. 2023). We focused on two asymmetric ground-truth regimes: a mutant-sparse/environment-dense regime (M sparsity = 0.2, *W* sparsity = 1.0) and a mutant-dense/environment-sparse regime (M sparsity = 1.0, *W* sparsity = 0.2). These settings plant reciprocal architectures at the same matrix dimensions as the empirical data, allowing us to ask whether the rotation diagnostic identifies the side of the map that is truly sparse. For each regime, two independently generated synthetic datasets were analyzed.

For each synthetic matrix, we applied the same SSD rotation-test procedure used for the empirical data. In the mutant-side test, the mutant axis was rotated by right-multiplying the fitness matrix by a random orthogonal matrix (*F′* = *FO_L_*), while fixing λ*_W_* = 10*^—^*^4^ and sweeping λ*_M_* over 25 logarithmically spaced values from 10*^—^*^3^ to 1.5. In the environment-side test, the environment axis was rotated by left-multiplying the fitness matrix by a random orthogonal matrix (*F′* = O*_E_F*), while fixing λ*_M_* = 10*^—^*^3^ and sweeping λ*_W_* over 25 logarithmically spaced values from 10*^—^*^4^ to 10*^—^*^2^. Two rotation replicates were used for each synthetic dataset and each mode. The plotted points in Supplementary Figure S15 are the mean original and rotated SSD solutions at each regularization setting, averaged across the two synthetic datasets. This calibration shows that sparse structure can be detected at the empirical matrix dimensions, but it also cautions against treating raw mutant-side and environment-side rotation gaps as directly comparable numerical measures of sparsity.

### 4.10 Visualizing loading matrices to interpret the meaning of each phenotypic axis (Figure 6)

To facilitate biological interpretation, we computed the loading matrices, which represent the contribution (weights) of each mutant and environment to these four abstract phenotypic dimensions. Specifically, mutant loadings were calculated by multiplying the mutant singular vectors (M) by the square root of the diagonal singular-value matrix (Σ), and environmental loadings were calculated by multiplying the environmental singular vectors (E) by the square root of Σ. These loading matrices allowed us to interpret each phenotypic dimension regarding underlying biological mechanisms related to drug resistance (Figure 6).

### 4.11 Data Availability Statement

All data and code are available at https://github.com/Russellrezagh/SVD-SSD-genotype-phenotype-mapp

## Acknowledgments

Funding for this work was provided by a National Institutes of Health grant R35GM133674 (to KGS), and a Alfred P Sloan Research Fellowship in Computational and Molecular Evolutionary Biology grant FG-2021-15705 (to KGS). We would also like to thank the members of the ASU Biodesign Center for Mechanisms of Evolution and the Geiler-Samerotte Lab for their support and feedback.

## Supplementary Material

**Figure S1:**
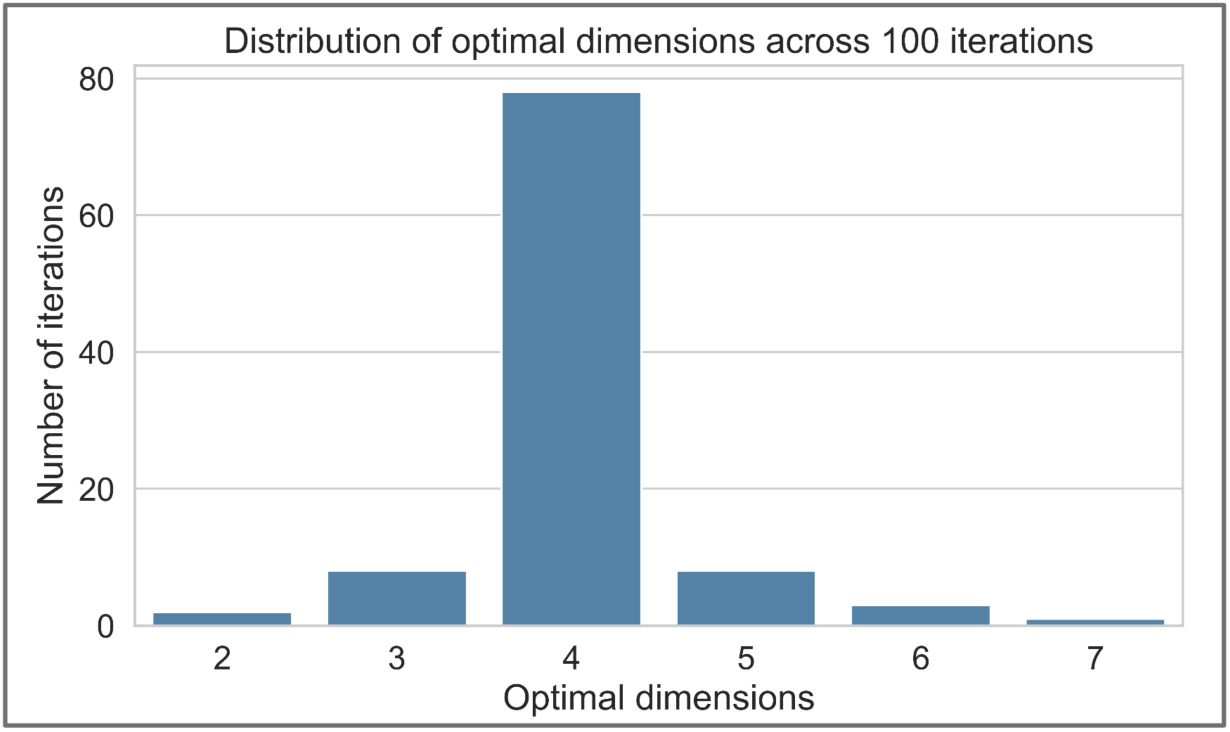
Distribution of optimal SVD dimensionality across repeated training iterations in Figure 2. For each of 100 repeated SVD runs related to the analysis in Figure 2, the model was trained using a different random subset of training mutants, and the optimal number of dimensions was selected for that iteration according to model performance. The histogram shows the frequency with which each dimensionality was chosen across iterations. In most runs, the optimal dimensionality was 4, indicating that the low-dimensional structure underlying the fitness matrix is stable across different training subsets.

**Figure S2:**
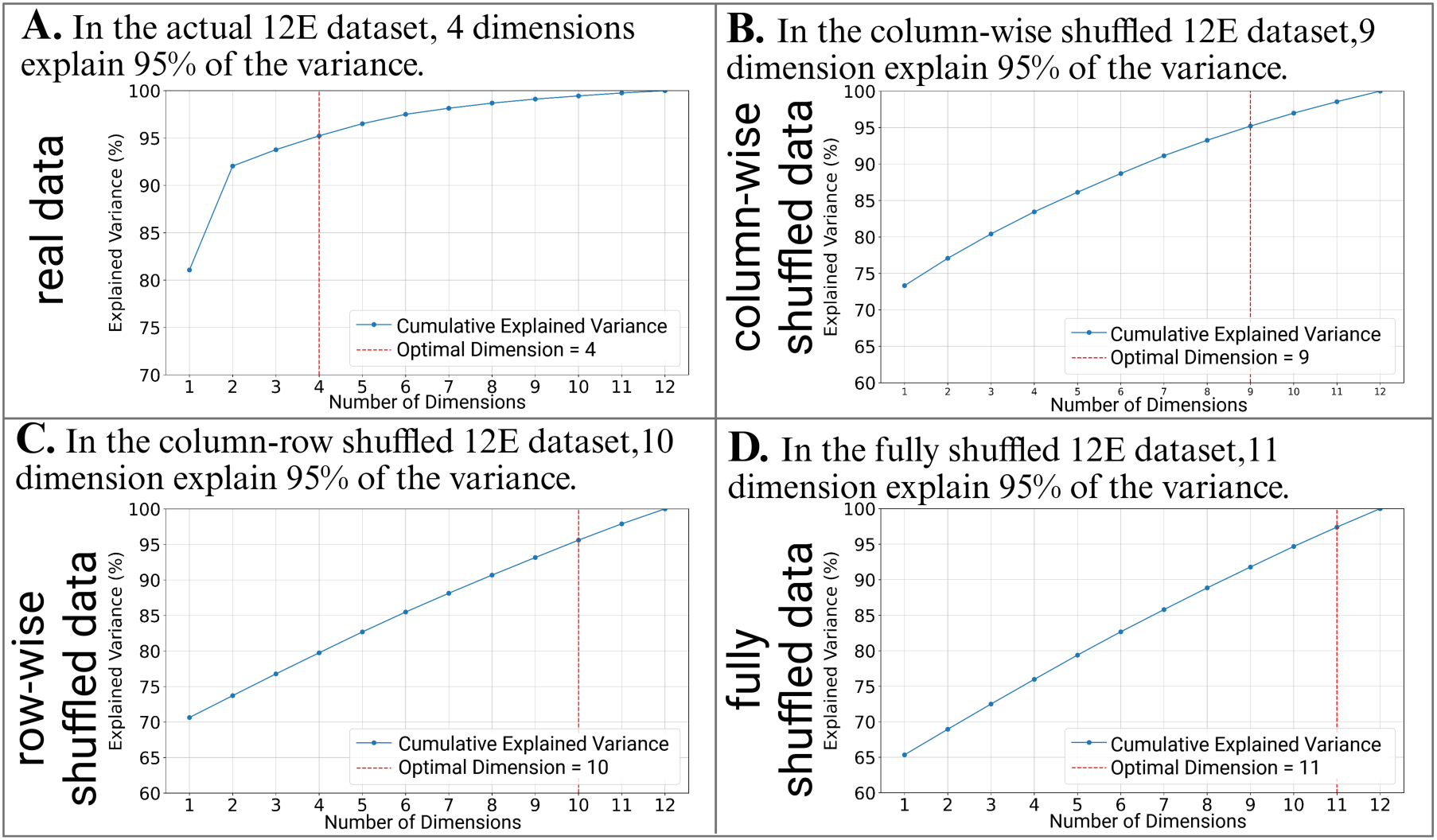
The fitness matrix we study contains stronger low-dimensional structure than shuffled controls. **(A)** Cumulative explained variance of the singular value decomposition (SVD) for the original 12-environment (12E) fitness matrix. In this focal dataset, 4 dimensions are sufficient to explain 95% of the total variance. **(B–D)** Cumulative explained variance for shuffled versions of the same 12E dataset, showing how many dimensions are required to explain the same amount of variance after disrupting matrix structure. **(B)** In the column-wise shuffled matrix, 9 dimensions are required to explain 95% of the variance. **(C)** In the row-wise shuffled matrix, 10 dimensions are required. **(D)** In the fully shuffled matrix, 11 dimensions are required. These results indicate that the original 12E fitness matrix contains substantial non-random low-dimensional structure that is lost after shuffling.

**Figure S3:**
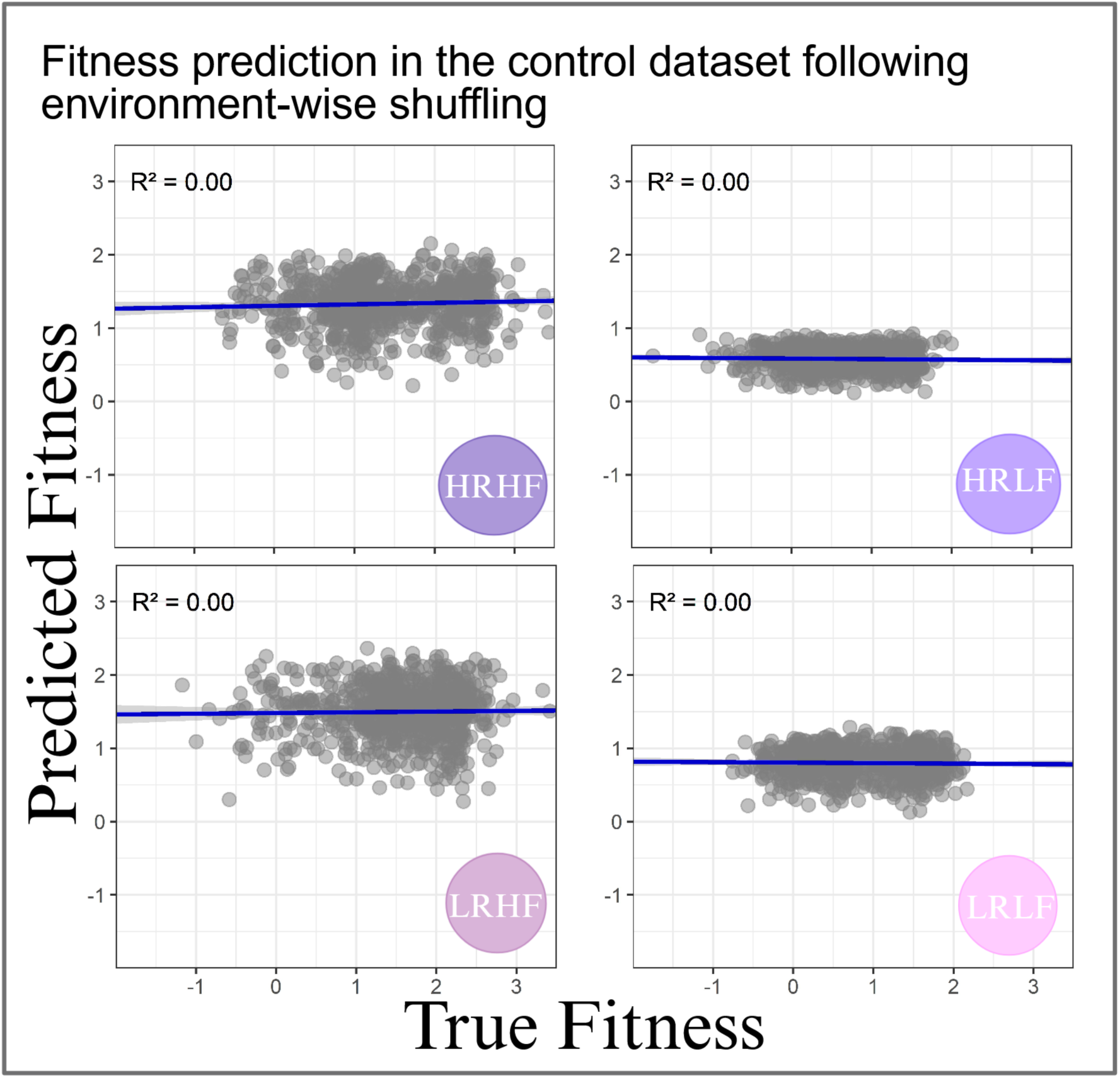
Double-drug prediction fails in a shuffled version of the 12E fitness matrix. Predicted versus true fitness in the four double-drug environments when the same SVD analysis used in Figure 2 was applied to a column-wise shuffled version of the full 12-environment (12E) fitness matrix. For this control, fitness values were randomly permuted across mutants independently within each environment, preserving the distribution of fitness values in each environment while destroying the correspondence of mutant identities across environments. The shuffled matrix was then split in the same way as described in Figure 2, using eight single-drug environments for training and four double-drug environments for testing. Each point represents the average prediction for each mutant, averaged across iterations. One hundred iterations in total were performed in each of which 100 mutations are randomly chosen as the training set, as in Figure 2. *R*^2^ values are shown in each panel. Prediction accuracy collapsed to approximately zero in all four double-drug environments, indicating that the predictive signal in the original analysis depends on structured mutant-specific relationships across environments.

**Figure S4:**
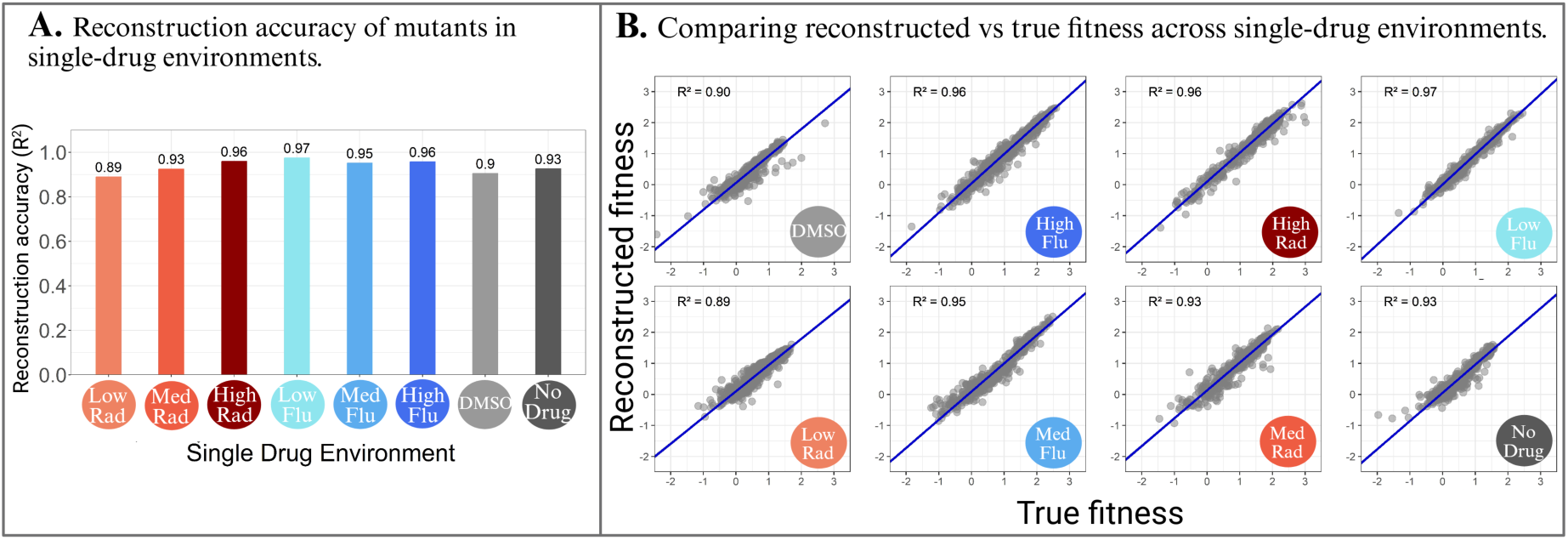
Reconstruction accuracy of the SVD model in the single-drug environments used for the training set. **(A)** Summary of the data in panel B. Barplots show the reconstruction accuracy (*R*^2^) for each of the eight single-drug environments used in the SVD framework shown in Figure2: three Radicicol concentrations (red; 5 **µ**M, 15 **µ**M, 20 **µ**M), three Fluconazole concentrations (blue; 4 **µ**g/ml, 6 **µ**g/ml, 8 **µ**g/ml), and two controls (gray; DMSO, no drug). **(B)** Scatter plots comparing reconstructed versus true fitness values for mutants in each single-drug environment. Each point represents one mutant. Reconstruction values were obtained by repeatedly running the model 100 times using random subsets of 100 mutants as the training set and estimating fitness for the remaining mutants; each mutant’s reported value is the average across iterations in which it was included in the training subset. These plots indicate that the low-dimensional SVD representation captures most of the structure in the single-drug fitness matrix that was used for the double-drug predictions in Figure 2.

**Figure S5:**
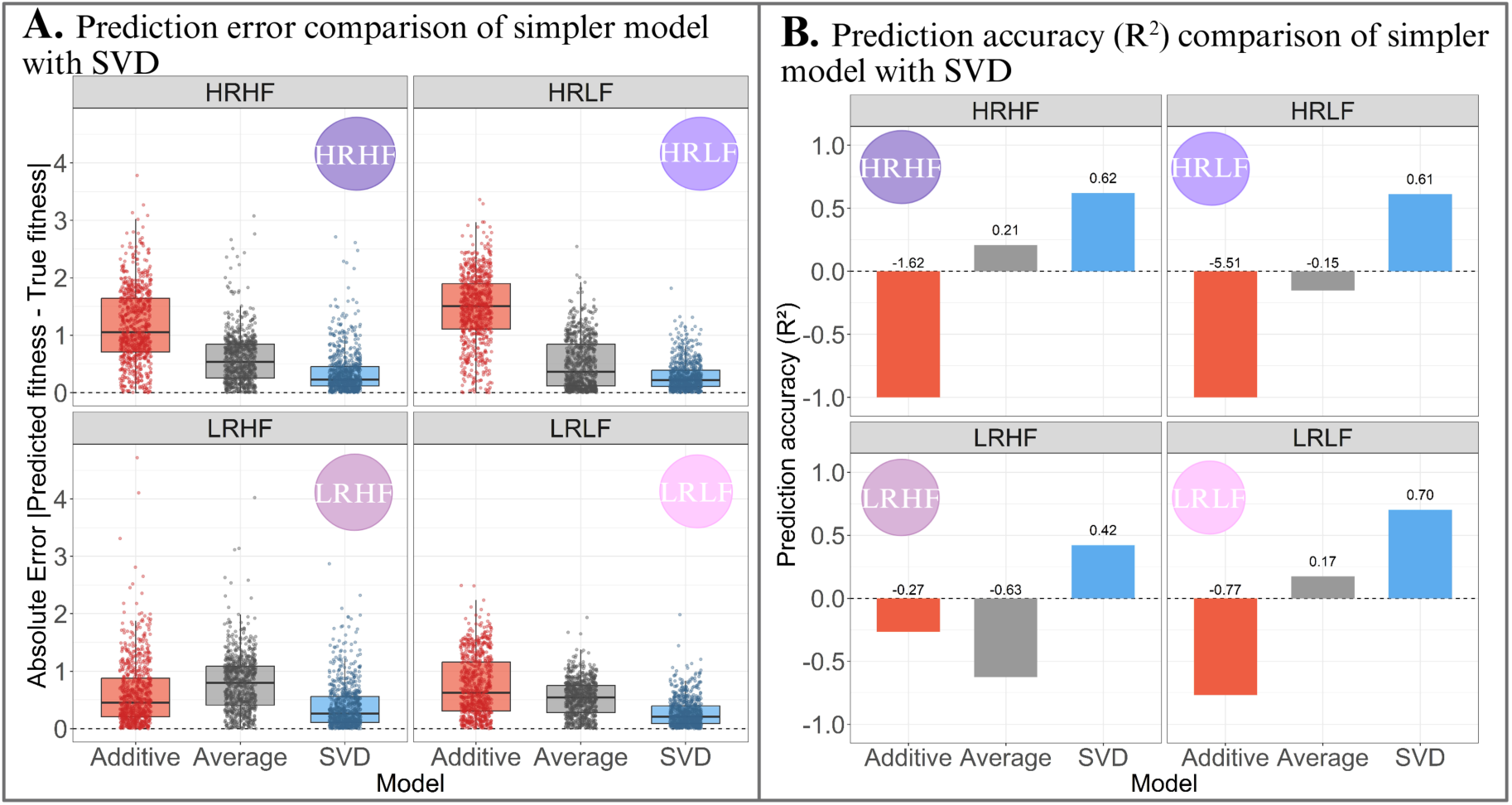
SVD fitness predictions outperform simple additive or average models. **(A)** Absolute prediction error for mutant fitness in the four double-drug environments (HRHF, HRLF, LRHF, LRLF) when using three models: an additive model (red), an average model (grey), and the SVD model (blue). Each point represents one mutant, and boxplots summarize the distribution of absolute errors (predicted fitness minus true fitness) in each double drug environment. The additive model predicts fitness in double drug conditions as the sum of each mutant’s fitness in the corresponding single drug conditions, while the average predicts each mutant’s fitness in double drug conditions as its average fitness across the two corresponding single drug conditions. **(B)** Prediction accuracy of the three models in the same four double-drug environments, measured by *R*^2^ between predicted and observed fitness values. The SVD model consistently outperformed the two simpler baseline models, indicating that the low-dimensional structure learned from single-drug environments captures predictive information not recovered by additive or averaging-based approaches.

**Figure S6:**
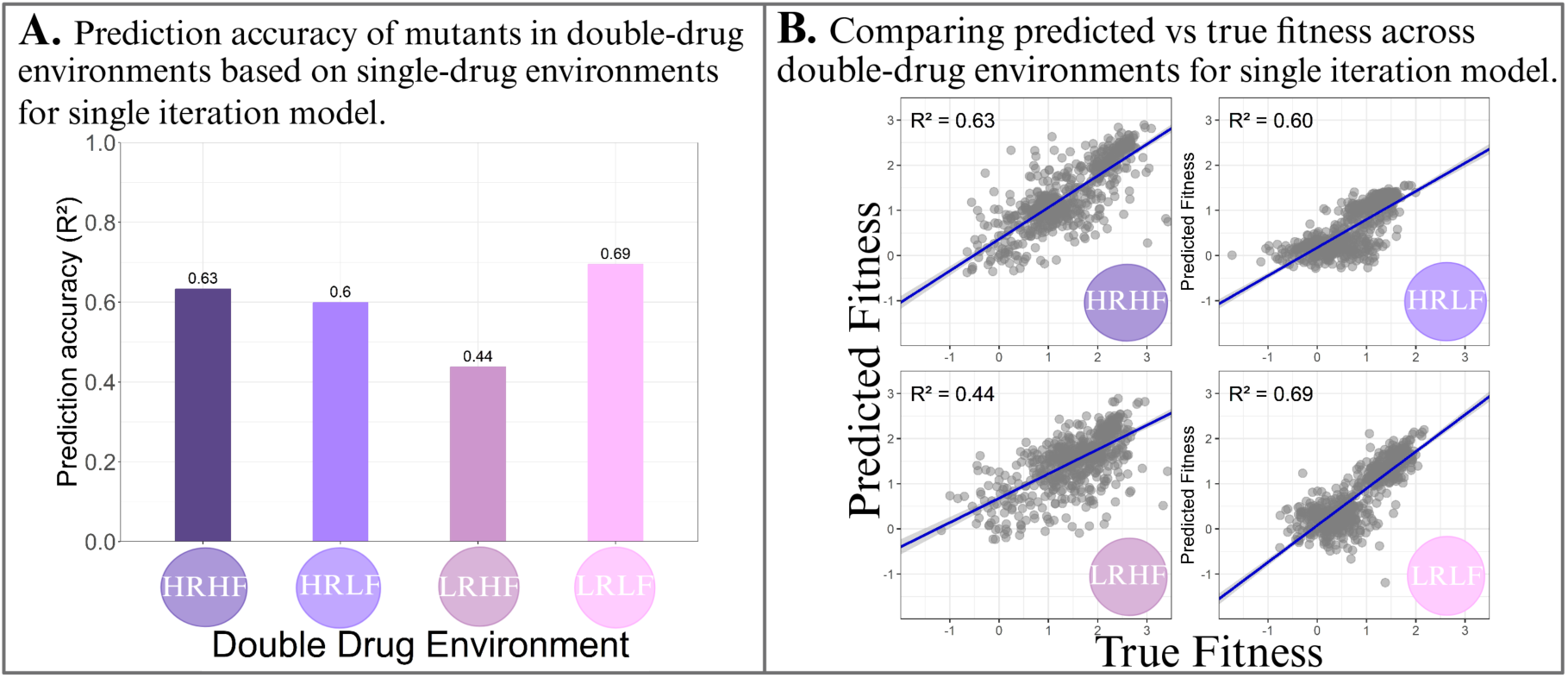
Prediction performance from a single SVD iteration is similar to the averaged results shown in Figure 2. **(A)** Prediction accuracy (*R*^2^) for the four double-drug environments obtained from one iteration of the SVD model, using the same training and testing framework as in Figure 2 but without averaging predictions across 100 iterations. **(B)** Predicted versus true fitness for 674 mutants in each double-drug environment for that run. Although prediction accuracy is slightly reduced relative to the 100-iteration average in Figure 2, the same overall pattern is preserved across environments, indicating that repeated averaging primarily improves robustness rather than creating the predictive signal.

**Figure S7:**
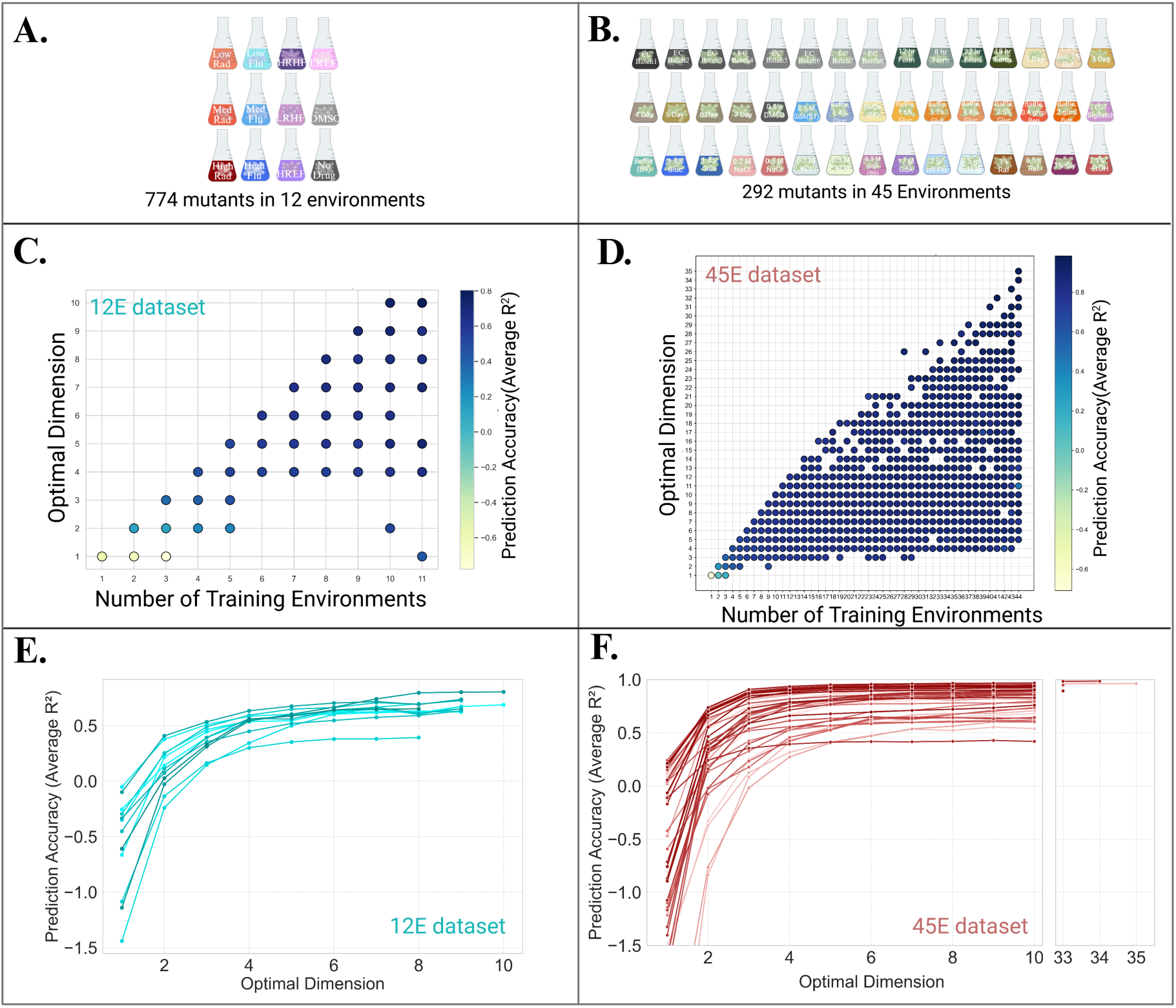
Prediction accuracy saturates with a small number of optimal dimensions and/or training environments. **(A,B)** Schematic summaries of the two datasets analyzed in Fig. 3 (A) The 12E dataset from (Schmidlin et al. 2024), containing fitness measurements for 774 mutants across 12 environments. **(B)** The 45E dataset from (Kinsler et al. 2020), containing fitness measurements for 292 mutants across 45 environments. **(C,D)** Prediction accuracy (average *R*^2^) from the Fig. 3 analysis, shown as a function of both the number of training environments and the optimal number of SVD dimensions. Each point corresponds to a model fit for a given number of training environments and particular optimal dimension, colored by average prediction accuracy on held-out environments. **(C)** 12E dataset. **(D)** 45E dataset. In both datasets, high prediction accuracy is achieved with relatively few training environments and a limited number of dimensions, while further increases provide only small additional gains. **(E,F)** Prediction accuracy from the Fig. 3 analysis plotted as a function of optimal dimensionality. Each line represents results from a different test environment. **(E)** 12E dataset. **(F)** 45E dataset. In both datasets, prediction accuracy increases rapidly with the first few dimensions and then plateaus, indicating that only a small number of phenotypic dimensions are needed to capture most of the predictive signal.

**Figure S8:**
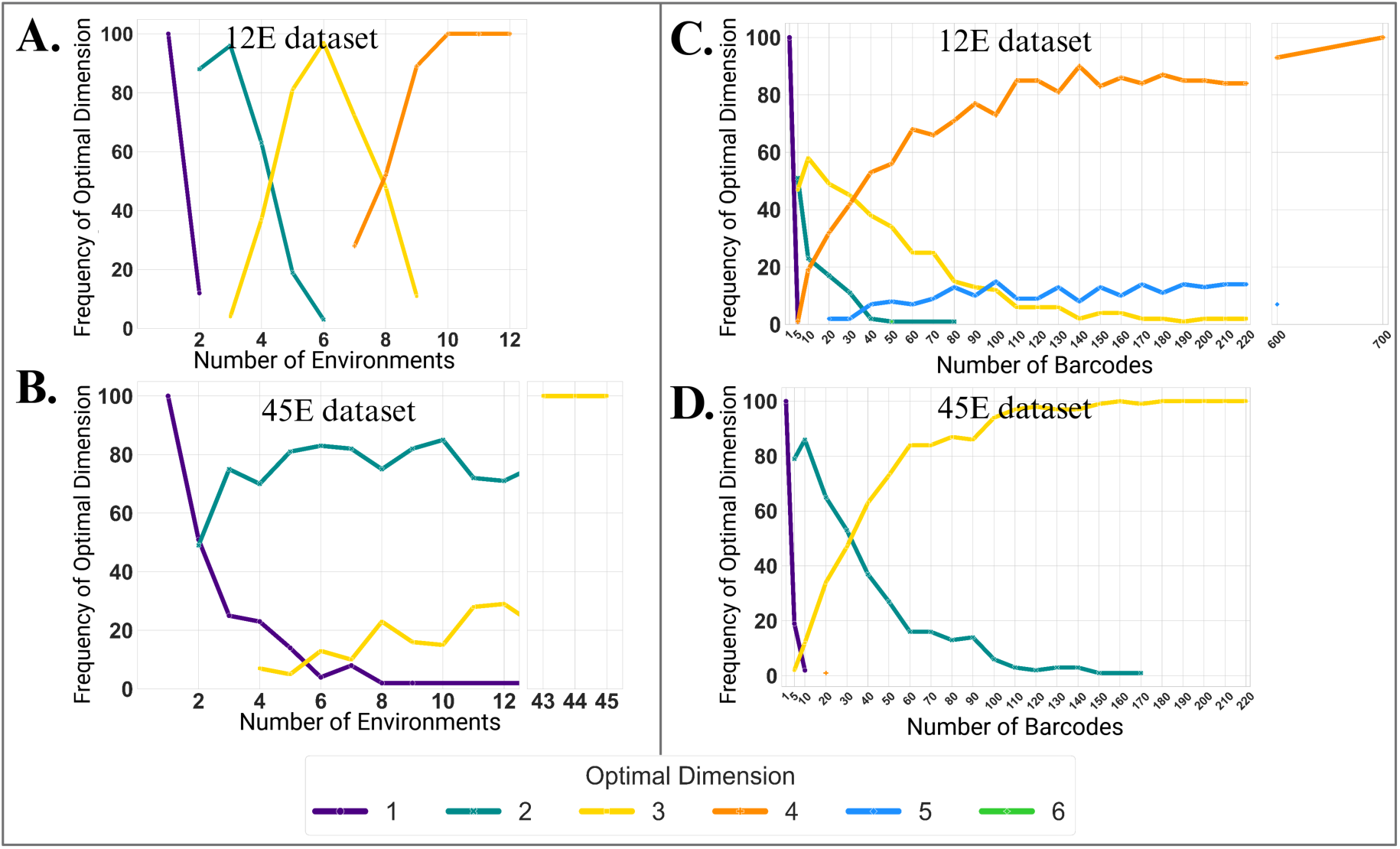
A dataset with 45 environments (Kinsler et al. 2020 (Kinsler et al. 2020); bottom) is captured in less latent dimensions than a dataset with only 12 environments (Schmidlin et al. 2024 (Schmidlin et al. 2024); top). This figure shows how the dimensionality of the fitness data changes when varying the number of environments (left) or the number of mutants (i.e. “barcode”) (right). For this analysis, the “optimal dimension” was defined as the minimum number of dimensions required to explain 95% of the variance in the training data. On the left, the training data includes all mutants, and the number of environments on the horizontal axis. On the right, the training data includes all environments, and the number of barcoded mutants on the horizontal axis. For each value on the horizontal axis, the SVD analysis was performed 100 times by iteratively subsampling environments (left) or mutants (right). The y-axis shows the frequency of each level of optimal dimensions as a function of either the number of environments sampled (left) or the number of mutants sampled (right). In both cases, the 12E dataset requires up to four dimensions to capture variation in fitness, while the 45E dataset, despite spanning more environments, stabilizes at two or three dimensions.

**Figure S9:**
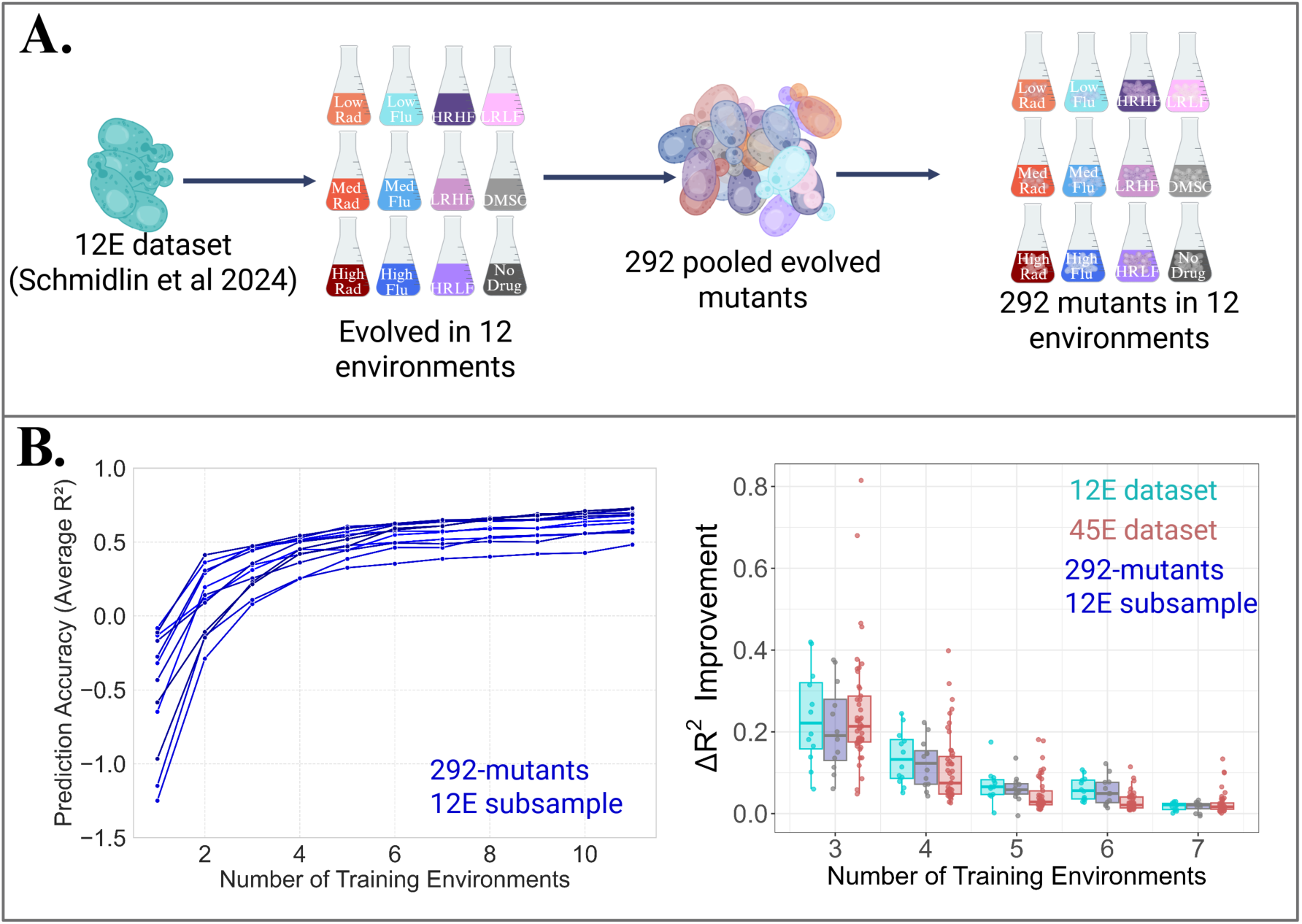
Reducing the 12E dataset to the same number of mutants (292) as the 45E dataset does not eliminate the difference in prediction-saturation behavior. **A** Schematic of the size-matched 12E subsample. From the original 12E dataset (Schmidlin et al., 2024), 292 mutants were selected to match the size of the 45E dataset. Mutants were sampled to preserve the relative proportions of the clusters defined in Schmidlin et al. (2024), so that the subsample retained the overall distributional structure of the full dataset. Fitness of the resulting 292-mutant 12E subsample was then analyzed across the same 12 environments. **B** Prediction behavior of the 292-mutant 12E subsample in the Fig. 3 analysis. Left, average prediction accuracy (*R*^2^) as a function of the number of training environments; each line corresponds to one held-out test environment. The size-matched 12E subsample still shows strong prediction performance and rapid saturation with increasing numbers of training environments. Right plot, incremental improvement in prediction accuracy (Δ*R*^2^) obtained by adding one additional training environment, compared across the full 12E dataset, the 45E dataset, and the 292-mutant 12E subsample. The Δ*R*^2^ pattern of the size-matched 12E subsample remains more similar to that of the full 12E dataset than to that of the 45E dataset, indicating that the faster saturation observed in the 45E dataset is not explained simply by the smaller number of mutants.

**Figure S10:**
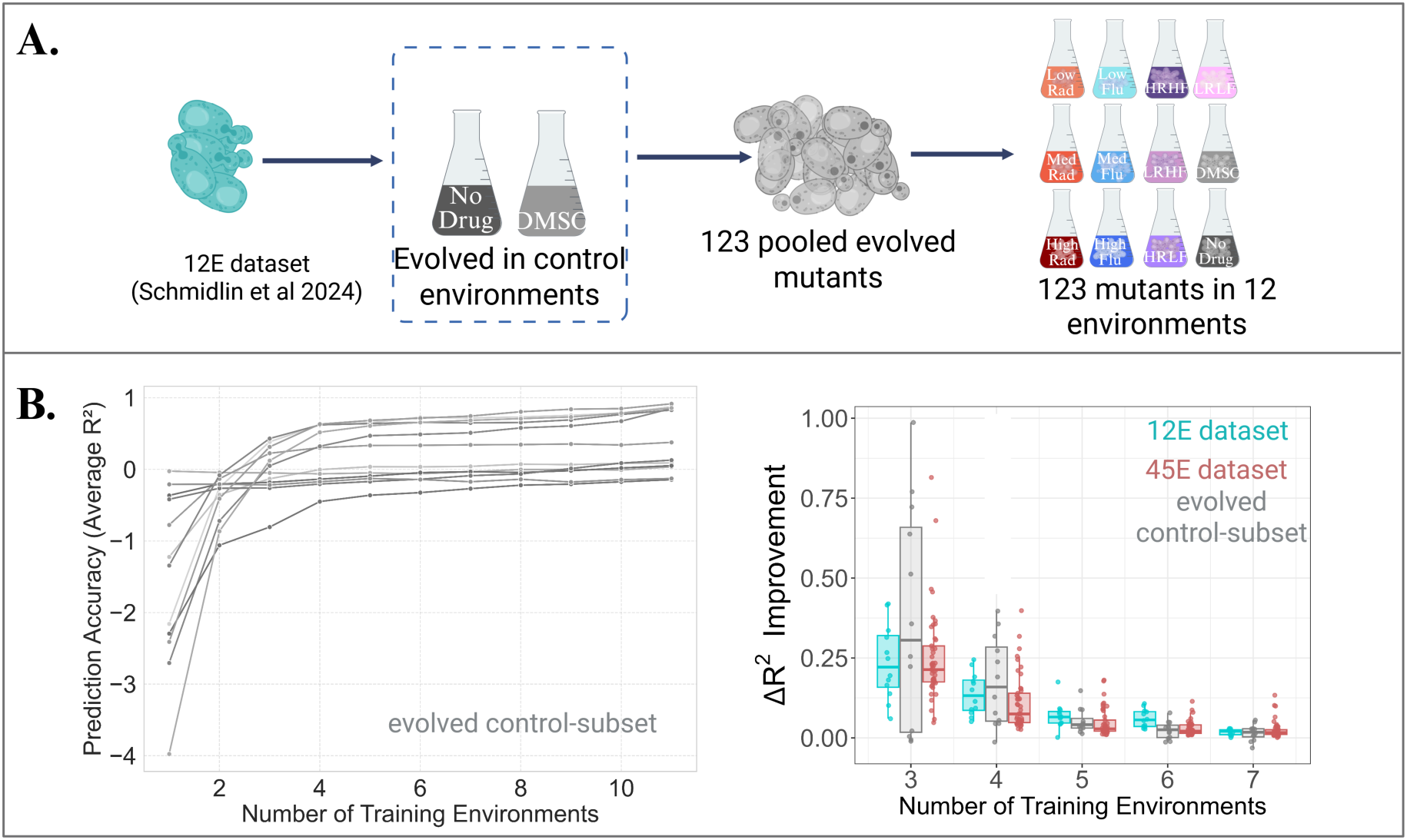
Down-sampling the 12E dataset such that it contains only mutations that evolved in a single selection pressure (glucose limitation) reveals prediction dynamics that are more similar to the 45E dataset. **A** Diagram of the analyzed subsample consisting of 123 mutants from the 12E dataset that evolved in the control environments only (‘DMSO’ and ‘No Drug’), with fitness measurements pertaining to all 12 assay environments. **B** For each number of training environments, 100 randomized iterations were performed, and prediction accuracy on held-out environments was averaged across iterations, as in Fig. 3C. The left plot indicates average *R*^2^ across test environments. The right plot indicates improvement in prediction accuracy (Δ*R*^2^) as additional training environments are included. Although prediction accuracy still rises rapidly and saturates after a small number of environments, the pattern of Δ*R*^2^ is more similar to that of the 45E dataset than to that of the full 12E dataset, especially around 5–6 training environments. This suggests that a more uniform evolutionary background could be associated with lower effective complexity in fitness prediction.

**Figure S11:**
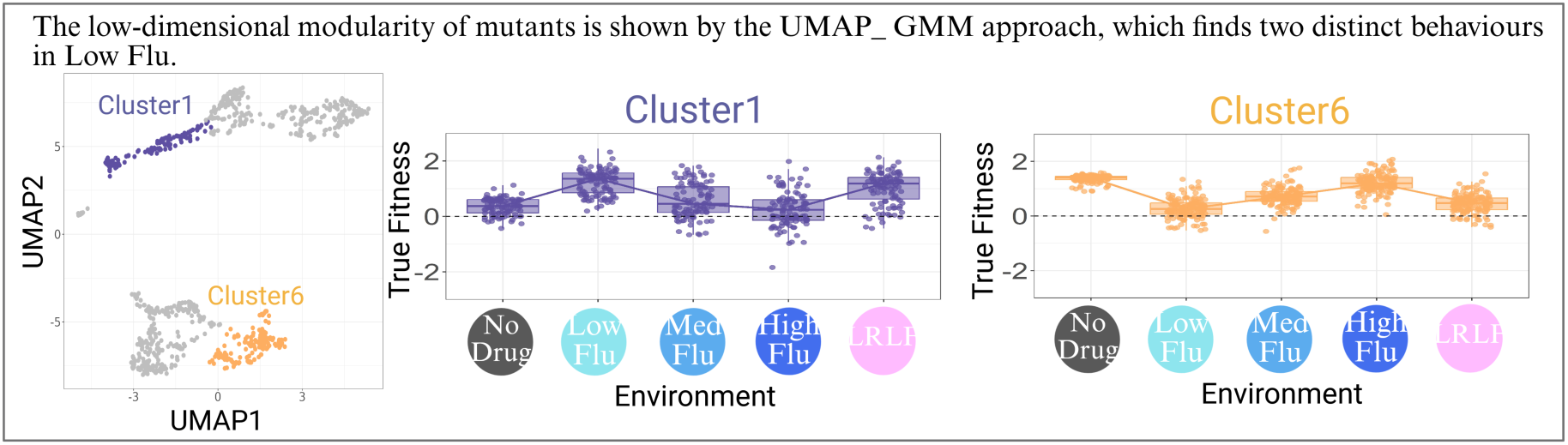
Mutant groups identified by prediction error in figure 4 correspond to clusters independently discovered in a previous study. To provide external validation for the novel mutant groups identified in Figure 4, we show that they correspond precisely to previously identified groupings (Schmidlin et al. 2024). These previous groupings were identified using an independent, unsupervised method (UMAP dimensionality reduction and Gaussian Mixture Model clustering) to categorize mutants based on their G x E interactions across the same 12 environments studied here. **Left:** The UMAP embedding of the groupings identified in previous work (Schmidlin et al. 2024), where the 774 drug-resistant mutants are represented as points. Here, two of the previously reported clusters are highlighted, including Cluster 1 (purple) and Cluster 6 (orange). **Middle:** Cluster 1 mutants display a characteristic non-monotonic fitness advantage with a distinct peak in the Low Fluconazole environment. This behavior mirrors that of the “most underpredicted” mutants identified through our prediction-error analysis (Figure 4C). **Right:** The fitness profile of mutants in Cluster 6 show a non-monotonic fitness decline in Low Fluconazole. This pattern is identical to that of the “most overpredicted” mutants identified by our method (Figure 4E).

**Figure S12:**
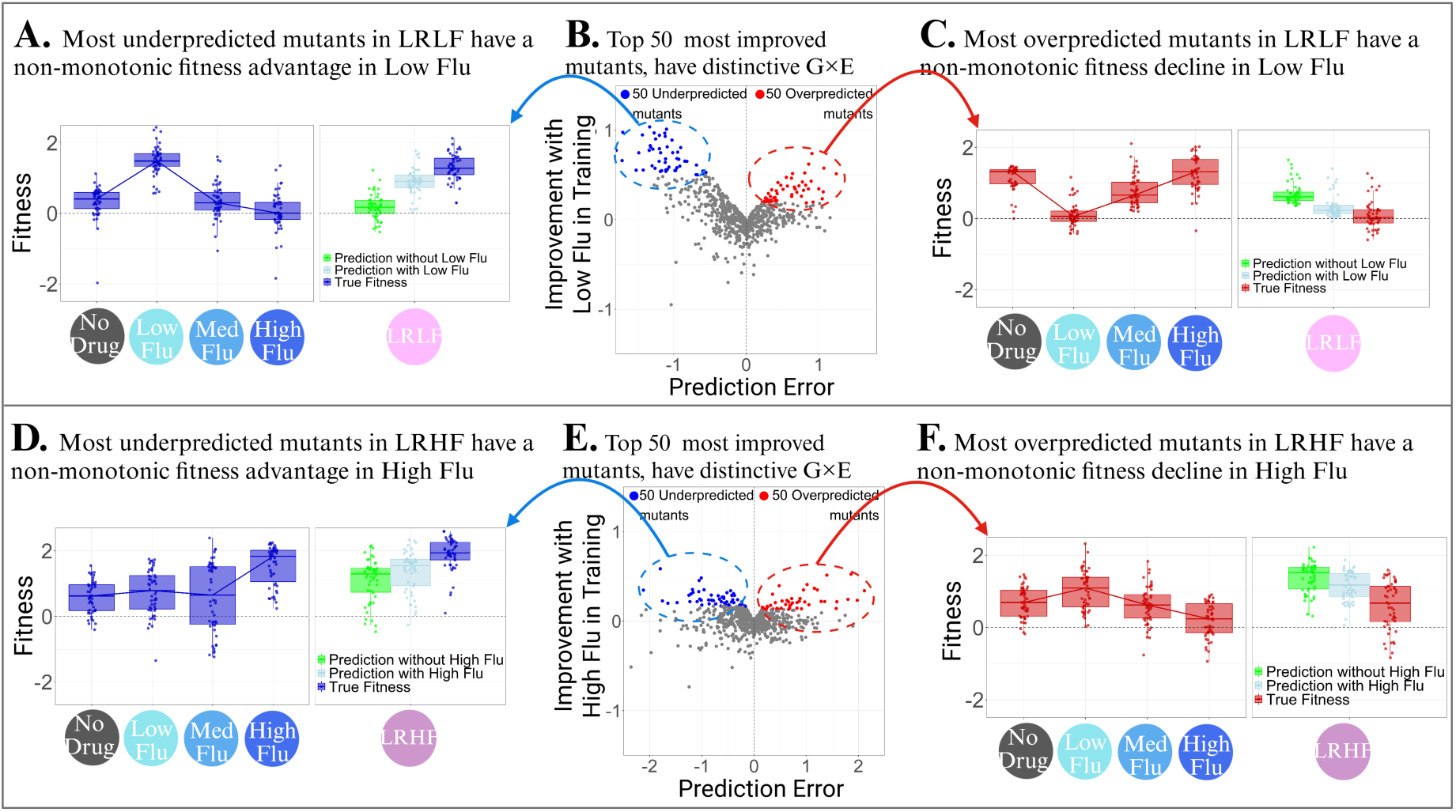
The mutants with fitness predictions that rely on data from the high flu environment have nonmonotonic changes in fitness across fluconazole concentrations. This figure extends the prediction-error analysis from figure 4 to investigate the LRHF environment, which is consistently difficult to predict. The top panel recapitulates figure 4 (LRLF) while the bottom panel pertains to the new LRHF analysis. The results suggest that some prediction difficulty in the LRHF environment is driven by nonmonotonic gene-by-environment (G⇥E) interactions across fluconazole, similar to those identified for the LRLF environment in figure 4. **(A–C)** Reproduction of figure 4. **(D)** Fitness profiles of the 50 most underpredicted mutants in LRHF as identified in panel E. These mutants display a non-monotonic fitness advantage that peaks in the High Fluconazole environment. The model trained without High Flu (green) is blind to this fitness peak, resulting in underprediction in LRHF. Adding High Flu in the training set improves the model’s approximation of fitness in LRHF (light Blue). **(E)** The relationship between prediction error when High Flu is excluded from training, and prediction improvement when High Flu is included. The 100 most improved underpredicted (blue) and overpredicted (red) mutants are highlighted. **(F)** Fitness profiles of the 50 most overpredicted mutants in LRHF as identified in panel E. These mutants exhibit a non-monotonic fitness relationship with fluconazole concentration, peaking in low fluconazole, and plummeting in high fluconazole. These mutants overlap strongly with those in panel A. The model misses the fitness decline in higher flu when it is excluded from the training and overpredicts fitness in LRHF (green). When High Flu is added to the training environment, the prediction improves (light Blue).

**Figure S13:**
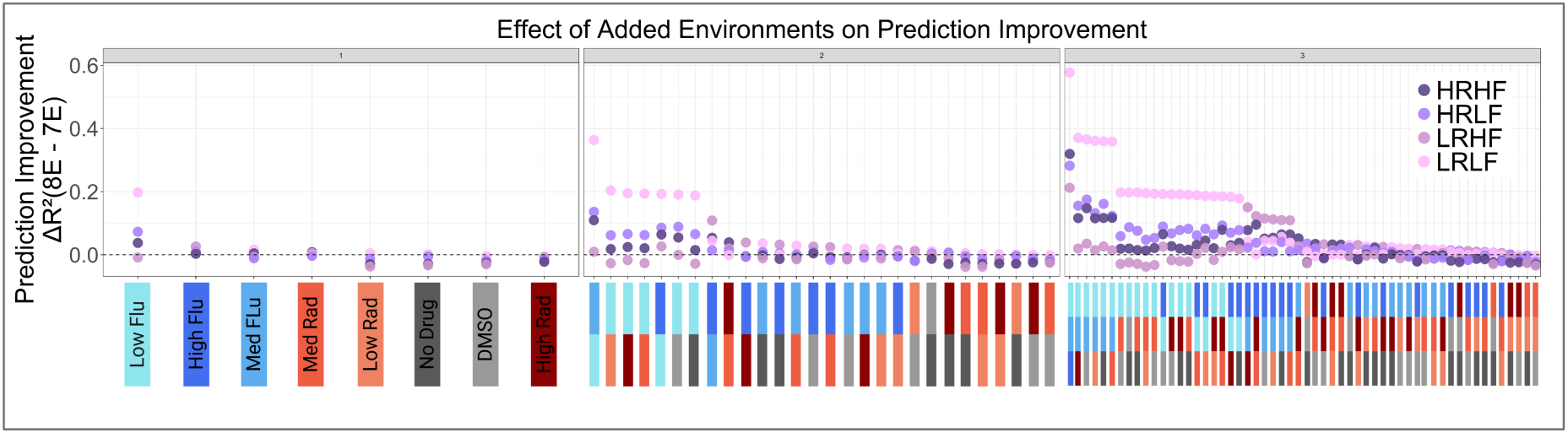
Low Fluconazole consistently provides the greatest prediction improvement regardless of training set composition. To robustly test the informational value of individual single-drug environments, this figure expands upon the analysis in Figure 4B. Here, we systematically hold out environments from the training set, and then report the improvement in prediction accuracy upon adding those training environments back. The y-axis shows the improvement in prediction accuracy (Δ*R*^2^). The colored bars at the bottom of each panel indicate the specific environment or combination of environments that was held out and then added back to the training set. Within each panel, the combinations are sorted from left to right by their impact on prediction improvement. **Left:** Shows the effect of adding each of the 8 single-drug environments individually to a 7-environment training set. As shown in Figure 4B, adding Low Fluconazole (light blue bar) provides the largest single improvement, especially for the LRLF Environment (light pink dot). **Middle:** Shows the effect of adding all unique pairs of environments to a 6-environment training set. The combinations yielding the highest prediction improvement (Δ*R*^2^) are consistently those that include Low Fluconazole. **Right:** Shows the effect of adding all unique triplets of single-drug environments to a 5-environment training set. Again, the most informative combinations are overwhelmingly those that contain Low Fluconazole.

**Figure S14:**
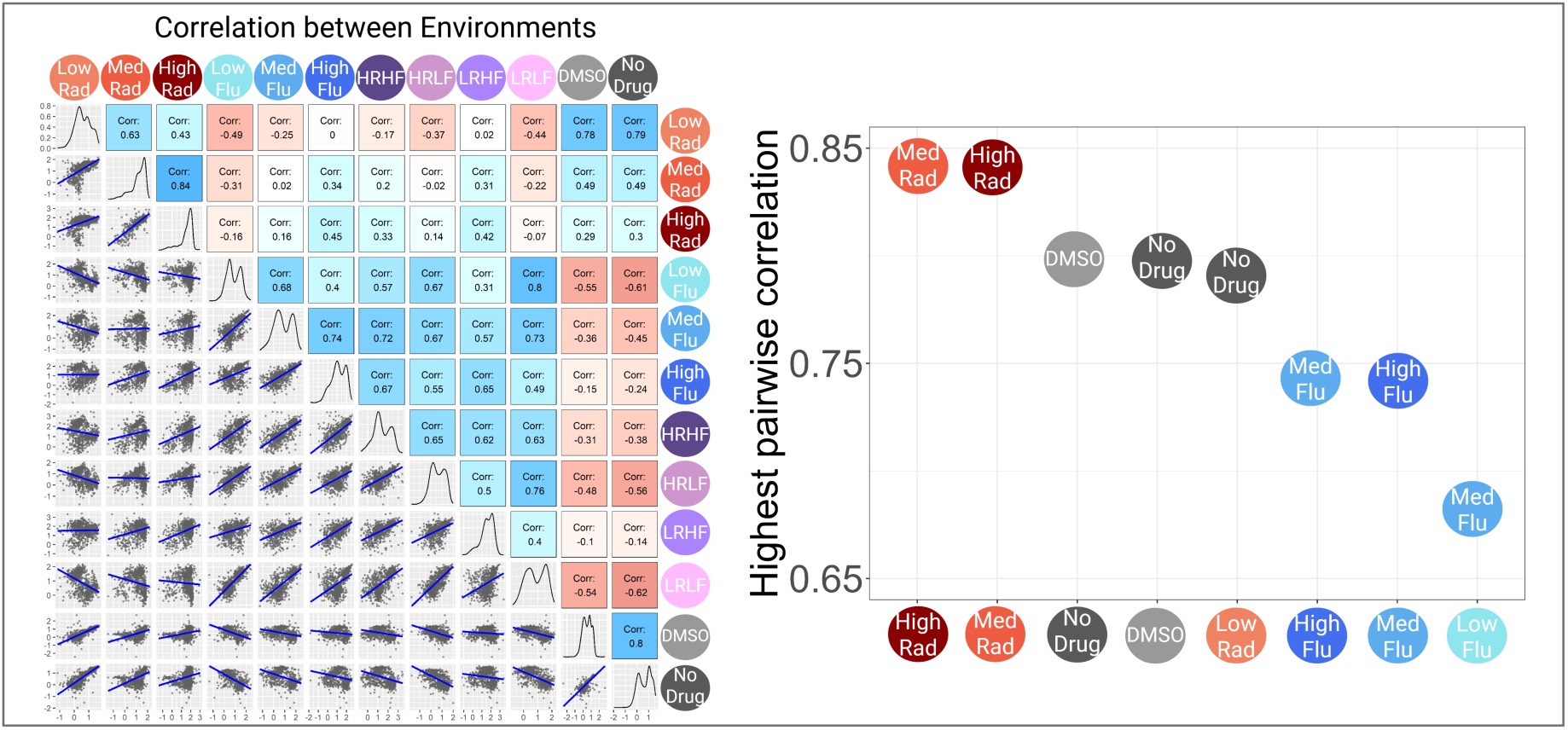
Low Fluconazole exhibits the least correlation with other single-drug environments. To further investigate why low-dose fluconazole is uniquely informative for predicting fitness, this figure analyzes the pairwise correlations of fitness between all 12 environments. **Left:** Shows a matrix of pairwise comparisons between all environments. The lower triangle contains scatter plots of fitness in each pair of environments, with a blue line showing the linear regression. The diagonal shows the density distribution of fitness values for each environment. The upper triangle displays the Pearson correlation coefficient for each pair, with the color indicating the strength and sign of the correlation, blue for positive, and red for negative. **Right:** Shows a summary plot displaying the highest pairwise correlation for each single-drug environment. For each environment on the x-axis, the y-value represents its maximum correlation with any other single-drug environment. This metric quantifies how redundant an environment is; a low value indicates high uniqueness. The analysis clearly shows that Low Fluconazole has the lowest maximum pairwise correlation (r = 0.68) compared to all other single-drug conditions. This indicates that the pattern of mutant fitness in Low Fluconazole is the most distinct and least redundant in the dataset.

**Figure S15:**
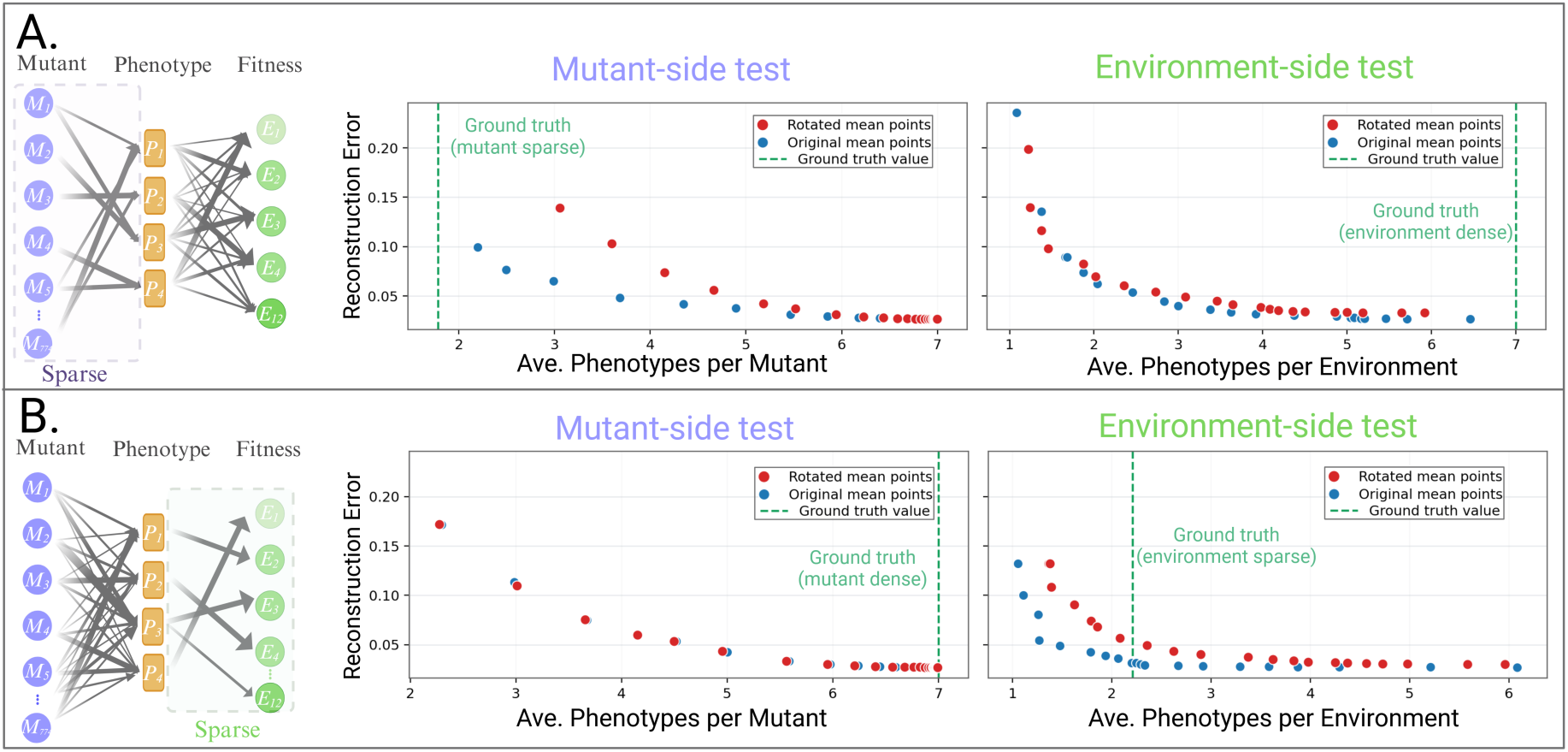
Synthetic calibration of the SSD rotation diagnostic under two asymmetric architectures. Synthetic fitness matrices were generated with 12 environments, 774 mutants, and 7 latent phenotypic dimensions. **(A)** Row A shows results for a dataset generated with asymmetric architecture in which the mutant-to-phenotype mapping is sparse (∽2 phenotypes per mutant) and the phenotype-to-environment mapping is dense (∽7 phenotypes per environment). **(B)** Row B shows the reciprocal architecture, in which the mutant-to-phenotype mapping is dense and the phenotype-to-environment mapping is sparse. In the schematic at left of each row, the dashed box marks the side of the map that was planted to be sparse. The two plots in each row compare SSD solutions for the original synthetic matrix (blue) and an orthogonally rotated control (red) for the mutant-side test and the environment-side test, respectively. Each point represents the mean solution at one regularization setting, plotted as reconstruction error versus inferred sparsity, measured as the average number of phenotypic dimensions per mutant or per environment. The green dashed vertical line marks the planted ground-truth sparsity value for that mode, and lower values on the horizontal axis indicate sparser solutions. This calibration tests the rotation diagnostic within each mode at the same dimensions as the empirical dataset, rather than treating raw mutant-side and environment-side gap sizes as directly comparable measures of sparsity.

**Figure S16:**
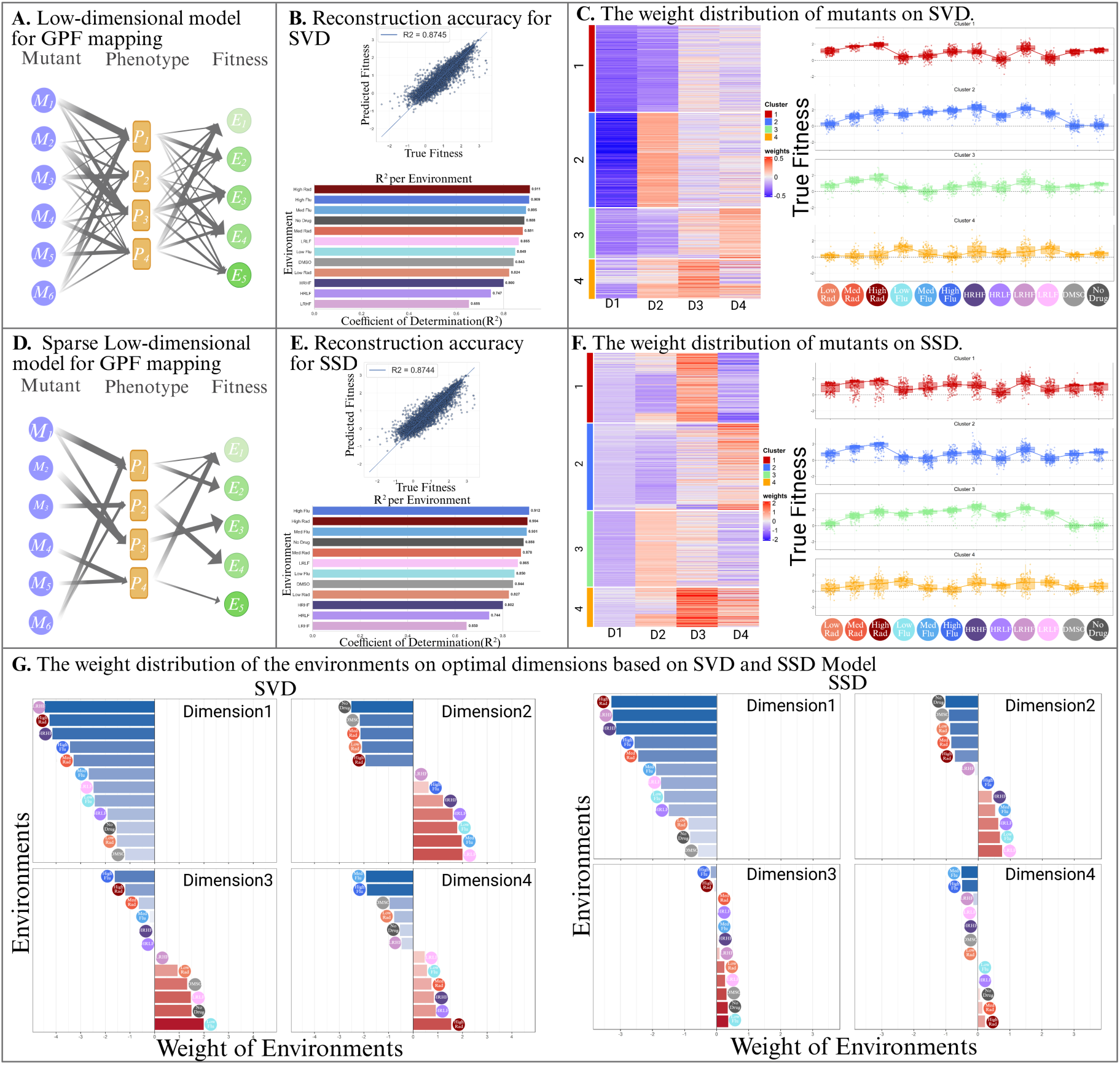
A sparse decomposition model (SSD) indicates a similar structure of the phenotypic dimensions identified by standard SVD. To assess the phenotypic dimensions identified in figure 6, this figure compares the standard Singular Value Decomposition (SVD) model with Sparse Structure Discovery (SSD). Both models were applied to the 12E fitness dataset (774 mutants x 12 environments) to interpret its underlying structure. Panels A to C indicate Standard SVD Model Analysis. **(A)** A conceptual schematic of a low-dimensional genotype-phenotype-fitness map inferred from SVD. **(B)** The model accurately reconstructs the original fitness values with an overall *R*^2^ = 0.8745; also, the bottom plot represents the reconstruction accuracy based on the environment. **(C)** A heatmap of mutant weights across the four phenotypic dimensions, with the corresponding fitness profiles for the resulting mutant clusters. Panels *D* to *F* show the parallel results obtained using the SSD algorithm. **(D)** A conceptual schematic of a low-dimensional genotype-phenotype-fitness map inferred from SSD. **(E)** The reconstruction accuracy of the SSD model with overall *R*^2^ = 0.8744 indicates that applying a sparsity constraint does not reduce the model’s ability to explain the variance in the data. **(F)** The corresponding heatmap of mutant weights and cluster fitness profiles from the SSD model. The fundamental structure and biological interpretation of the clusters are similar to the SVD results. **(G)** Comparison of Environmental Weights based on SVD and SSD. This panel directly contrasts the environmental weights for each of the four dimensions as calculated by standard SVD on the left side and the SSD model on the right side. The analysis reveals that the core structure of each dimension, which includes the weights’ order and sign of the environments, is similar between the two methods.

